# Peeling Back the Layers: First Phylogenomic Insights into the Ledebouriinae (Scilloideae, Asparagaceae)

**DOI:** 10.1101/2020.11.02.365718

**Authors:** Cody Coyotee Howard, Andrew A. Crowl, Timothy S. Harvey, Nico Cellinese

**Affiliations:** Florida Museum of Natural History, University of Florida, Gainesville, FL 32611, USA; Department of Biology, University of Florida, Gainesville, FL 32611, USA; Duke University, Durham, NC 27708, USA; Plantae Novae, Thousand Oaks, CA 91360, USA; Biodiversity Institute, University of Florida, Gainesville, FL 32611, USA

**Author notes:** Corresponding author: Cody Coyotee Howard, Department of Biology, University of Florida, Gainesville, FL 32611, USA.

**Keywords:** Asparagaceae, *Drimiopsis*, *Ledebouria*, monocots, *Resnova*, Scilloideae

## Abstract

The Ledebouriinae (Scilloideae, Asparagaceae) are a widespread group of bulbous geophytes found predominantly throughout seasonal climates in sub-Saharan Africa, with a handful of taxa in Madagascar, the Middle East, India, and Sri Lanka. Phylogenetic relationships within the group have been historically difficult to elucidate. Here, we provide the first phylogenomic perspective into the Ledebouriinae. Using the Angiosperms353 targeted enrichment probe set, we consistently recovered four major clades (i.e., two *Ledebouria* clades, *Drimiopsis*, and *Resnova*). The two *Ledebouria* clades closely align with geography, either consisting almost entirely of sub-Saharan African taxa (*Ledebouria* Clade A), or East African and non-African taxa (*Ledebouria* Clade B). Our results suggest that the Ledebouriinae likely underwent a rapid radiation leading to rampant incomplete lineage sorting. We additionally find evidence for potential historical hybridization between *Drimiopsis* and a subclade within *Ledebouria* Clade A.

## 1. Introduction

Africa houses an enormous diversity of plants found across a range of habitats from tropical rainforests to arid landscapes (Couvreur et al., 2020; Linder, 2014). Still, there remain numerous, understudied African taxa that may refine hypotheses regarding evolution within Africa (e.g., the Rand Flora (Pokorny et al., 2015), the “arid track” (Balinsky, 1962), or showcase their own distinct patterns. One such group is the hyacinths (Scilloideae, Asparagaceae; formerly Hyacinthaceae; APG IV (The Angiosperm Phylogeny Group, 2016)), which consist of approximately 1,000 bulbous geophytes found throughout seasonal climates in Africa, as well as Madagascar, Europe, the Middle East, India, Sri Lanka, and South America (Speta, 1998). Digging into the evolution of the Scilloideae has improved our understanding of historical biogeography in Africa (Ali et al., 2013; Buerki et al., 2012), polyploidy (Jang et al., 2018), and ecology (Johnson et al., 2001; Vogel and Müller-Doblies, 2011). Unfortunately, the group overall remains largely understudied. To date, many clades within the Scilloideae have received varying degrees of phylogenetic and systematic attention (e.g., Ledebouriinae, Massonieae, Ornithogaloideae; (Lebatha et al., 2006; Martínez-Azorín et al., 2011; Pfosser et al., 2003; Venter, 2008); among many others), but most have been limited in taxonomic and/or geographic coverage. Large phylogenomic applications are almost nonexistent (Steele et al., 2012). To showcase the value of focusing attention on understudied groups such as the Scilloideae, we investigate the phylogenomic space of one of its subclades—the Ledebouriinae—the evolutionary relationships of which have been historically difficult to reconstruct.

The Ledebouriinae are widespread across sub-Saharan Africa, found predominantly within the “arid track” (Balinsky, 1962), with a handful of taxa in Madagascar, Yemen, India and Sri Lanka (Fig. 1a; (Giranje and Nandikar, 2016; Venter, 1993). The current center of diversity is the Limpopo, Mpumalanga, and KwaZulu-Natal regions of South Africa (Venter, 1993), with a secondary center in East Africa (Lebatha et al., 2006; Lebatha, 2004), both of which may simply reflect a geographic bias in our understanding of Ledebouriinae diversity. For example, throughout the distribution of the Ledebouriinae, studies continue to reveal undescribed diversity within the group (Howard, 2014; Ramana et al., 2012). Taxonomically, the group currently consists of *Ledebouria* Roth with two sections: *Drimiopsis* (Lindl. & Paxton) J.C. Manning & Goldblatt and *Resnova* (Van der Merwe) J.C. Manning & Goldblatt (Manning and Goldblatt, 2012; Manning, 2020; Manning et al., 2003). However, taxonomic concepts still vary and a lack of consistency in the hypotheses generated by phylogenetic studies has resulted in disagreements regarding the nature of the Ledebouriinae and whether this group includes one or more distinct entities (Lebatha et al., 2006; Manning et al., 2003). Historically, *Ledebouria* and *Drimiopsis* Lindl. & Paxton were classified as separate taxa, and *Drimiopsis* enjoyed its own identity for some time (Goldblatt et al., 2012; Lebatha et al., 2006; Müller-Doblies and Müller-Doblies, 2008). The status of *Resnova* van der Merwe, on the other hand, has always been in question (Goldblatt et al., 2012; Lebatha et al., 2006; Manning, 2020; Manning et al., 2003). Distinguishing characteristics of the three taxa at the generic level are few, but differences in floral morphology have been and remain to be the main characters used when determining the identity of a taxon. *Ledebouria* flowers often display fully reflexed tepals that surround a stipitate ovary, with a stem subtending the ovary and attaching it to the remaining floral structure. *Drimiopsis* flowers have connate, dimorphic tepals that often result in the flower having a closed appearance, and they lack a stipitate ovary (Hankey, 2003). *Resnova* ovaries also lack a stem, and flowers tend to have intermediate tepal morphology between *Ledebouria* and *Drimiopsis*, resulting in an often bell-shaped (campanulate) flower appearance. Leaf and bulb characters display considerable variation and have thus far been unreliable for use in identification of *Ledebouria, Drimiopsis* and *Resnova* (Lebatha et al., 2006), but have some significance for identifying a specimen within each taxon (i.e., distinguishing between species) (e.g., Fig. 2) (Lebatha, 2004; Manning, 2020; Venter, 1993). Currently, there are 64 *Ledebouria*, 14 *Drimiopsis*, and six *Resnova* species accepted (POWO, 2019). Phylogenetically, difficulties in reconstructing the relationships within the Ledebouriinae have stemmed from morphological homoplasy (e.g., leaf maculation) (Lebatha et al., 2006), the use of a small number of molecular markers with low information content (Lebatha et al., 2006; Manning et al., 2003), the lack of robust phylogenetic methods (Pfosser et al., 2003; Wetschnig et al., 2007), and the inclusion of a small fraction of the taxonomic and geographic diversity of the group. These shortcomings have resulted in various conclusions regarding the monophyly of taxa, ranging from debating the monophyly of each lineage (monophyletic; Lebatha et al. 2006, or polyphyletic;(Pfosser et al., 2003) to the taxonomic lumping of each lineage into a broadly circumscribed *Ledebouria* (Manning et al., 2003). These limitations also constrain our ability to detect confounding factors that have likely impacted the evolutionary history of the Ledebouriinae, which may be why it has been difficult to confidently reconstruct it.

**Figure 1.**
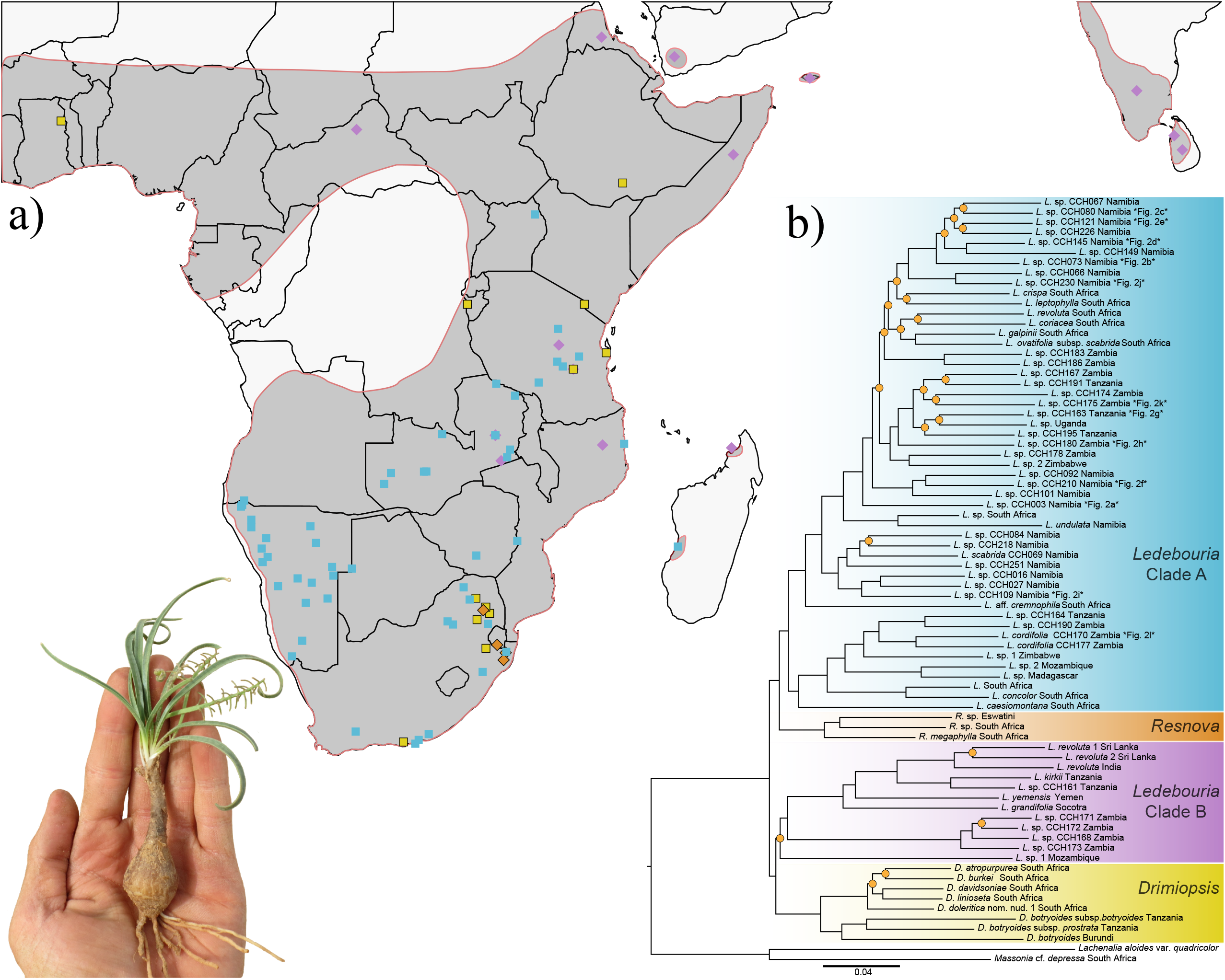
a) Localities of all samples included in this study (full dataset) overlaid on the general distribution of the Ledebouriinae (dark gray polygon). Shapes and colors correspond to clades shown in (b): *Ledebouria* Clade A (blue squares), *Ledebouria* clade B (purple diamonds), *Resnova* (outlined orange diamonds), *Drimiopsis* (outlined yellow squares). b) Maximum likelihood phylogenetic reconstruction of the Taxa70 dataset with the four major clades labeled. Orange circles indicate nodes with SH-aLRT and ultrafast bootstrap values *below* 80 and 95, respectively. Tips with a corresponding image in Fig. 2 are labeled.

**Figure 2.**
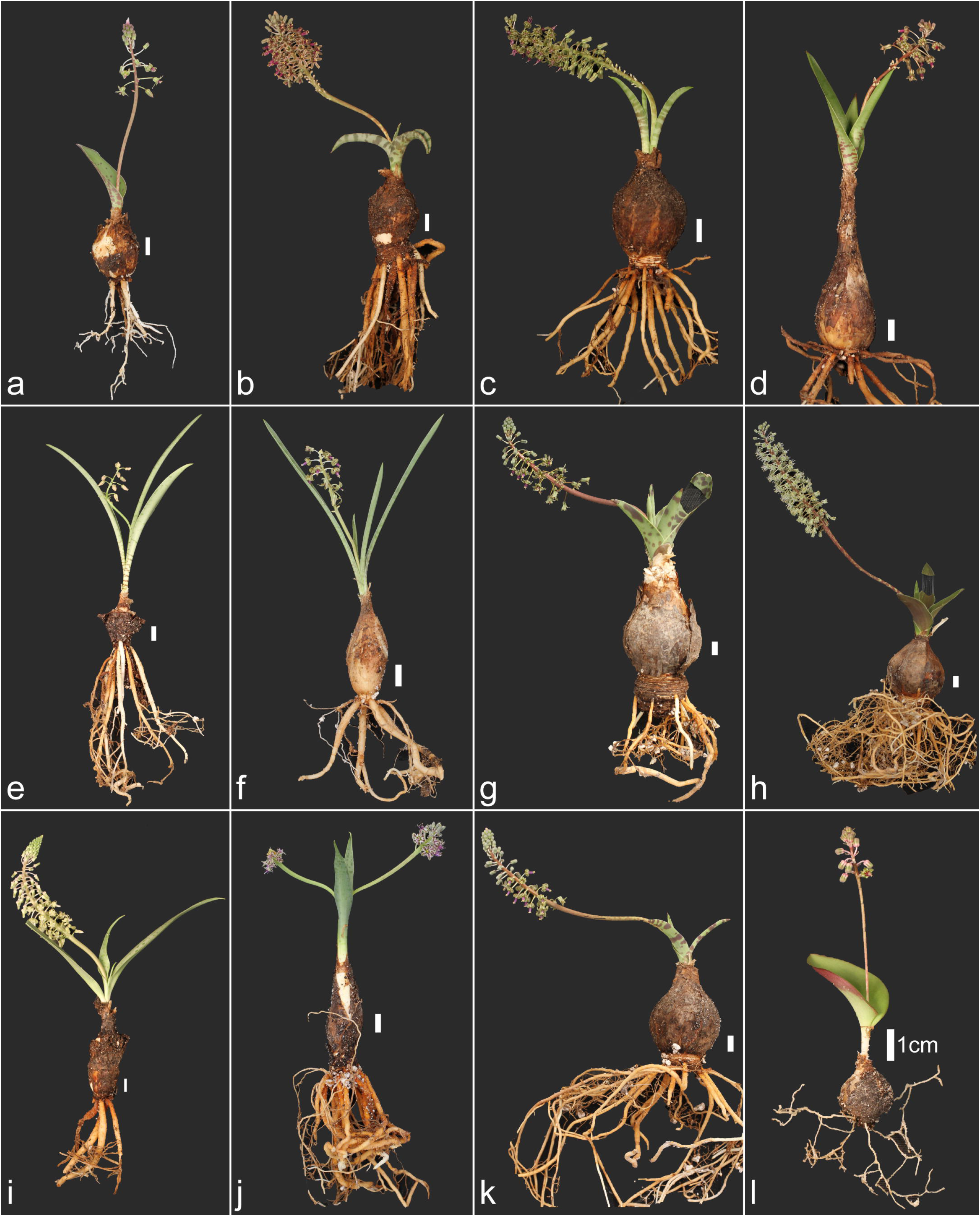
Examples of 12 *Ledebouria* species included in the phylogeny that are currently undetermined or undescribed, except for *L. cordifolia* (image l). These specimens showcase some of the range in morphology within the group, especially floral. Numerous attempts have been made to identify these specimens, but current species descriptions do not fit well. See Fig. 1 for each specimen’s placement within the phylogeny. White bars next to each specimen indicate a 1cm scale for that image. a) *Ledebouria* sp. CCH003 Namibia; b) *Ledebouria* sp. CCH073 Namibia; c) *Ledebouria* sp. CCH080 Namibia; d) *Ledebouria* sp. CCH145 Namibia; e) *Ledebouria* sp. CCH121 Namibia; f) *Ledebouria* sp. CCH210 Namibia; g) *Ledebouria* sp. CCH163 Tanzania; h) *Ledebouria* sp. CCH180 Zambia; i) *Ledebouria* sp. CCH109 Namibia; j) *Ledebouria* sp. CCH230 Namibia; k) *Ledebouria* sp. CCH175 Zambia; l) *Ledebouria cordifolia* CCH170 Zambia.

Based on previous dating analyses (Ali et al., 2012; Buerki et al., 2012), we hypothesize that the Ledebouriinae have experienced a rapid radiation(s). These studies suggest that the group originated during a time (i.e., within the last 30 My) when Africa was undergoing major climatic and geological shifts (e.g., increasing seasonality, orogenic activity) (Bobe, 2006; Jacobs, 2004; Partridge and Maud, 1987), and the bulbous geophytic habit may have allowed the Ledebouriinae to capitalize on such conditions (Howard et al., 2019). Additionally, hybridization and polyploidy within and between groups has been documented (Lebatha et al., 2006; Stedje and Nordal, 1987), which present further complications in reconstructing the evolutionary history of the group. Therefore, to recover a well-supported phylogenetic hypothesis for the Ledebouriinae, the use of a large, genomic dataset consisting of rapidly evolving markers is likely needed. Unfortunately, the group lacks the necessary genomic resources to develop a custom probe set. However, broadly applicable targeted-capture kits (Breinholt et al., 2021; Johnson et al., 2019) have been instrumental in bringing understudied groups, like the Ledebouriinae, to the phylogenomic playing field (Dodsworth et al., 2019).

Here, we provide the first phylogenomic insights into the Ledebouriinae by applying a HybSeq/targeted enrichment approach using the Angiosperms353 probe set, which has been effective in uncovering relationships at both deep and shallow scales (Larridon et al., 2019). The use of this probe set circumvents the lack of genomic resources that plagues the Ledebouriinae and allows future studies to leverage our dataset, which will enable continual refinement of the phylogenetic understanding of this group as well as the Scilloideae. We include samples from across the distribution of the group using both field-collected and museum-based specimens to investigate putative lineages within the Ledebouriinae (i.e., *Ledebouria, Drimiopsis* and *Resnova*) as well as potential reasons for discordance.

## 2. Materials and methods

### 2.1 Taxon sampling

Plants of Ledebouriinae were collected from the field in 2012 (Namibia), 2014 (Namibia), and 2017 (Tanzania, Zambia, and Namibia). Of the described Ledebouriinae species, we included 19 *Ledebouria*, six *Drimiopsis*, and three *Resnova*. Many *Ledebouria* samples included in our analyses are undescribed and are actively being studied for taxonomic description; similarly, we were unable to determine the status of other collections due to a lack of representative samples (e.g., they have only been collected at one site) coupled with a lack of flower morphology (i.e., plants have yet to flower ex situ), but their geographic distribution or unique leaf and/or bulb morphology warranted inclusion. Leaves of each sample were dried in silica gel for DNA extractions. Herbarium material from four institutions (Missouri Botanical Gardens, Uppsala’s Evolutionsmuseet Botanik, Sweden Museum of Natural History, and The Royal Botanic Gardens, Kew, DNA Bank, https://www.kew.org/data/dnaBank/) were also included.

Additionally, silica-dried leaf material from specimens with known provenance were generously donated from private collections. Our final dataset included 94 Ledebouriinae samples from across the distribution of the group as well as two outgroup taxa (*Massonia* cf. *depressa* Houtt. and *Lachenalia aloides* var. *quadricolor* (Jacq.) Engl.). See Table S1 for the specimen list with associated collection and locality information.

### 2.2 DNA extraction and probe selection

DNA extraction was carried out using a modified CTAB approach for all samples (Cullings, 1992; Doyle and Doyle, 1987). Extractions were quantified in a Qubit 2.0 (Invitrogen, Carlsbad, California, USA) using Qubit dsDNA Broad Range Assay Kit (Cat#: Q32850) following the manufacturer’s protocol. DNA concentrations were standardized between 30 – 75 ng/uL either by diluting solutions with deionized water or performing additional DNA extractions for later concentration. Afterwards, to ensure DNA was present, 37 random samples were visualized on a 2% agarose gel. Approximately 50 uL of each sample was sent to RAPiD Genomics (Gainesville, Florida, USA) for library preparation and DNA sequencing on an Illumina HiSeq 2000 sequencer (Illumina, San Diego, California, USA) using 2 x 150 bp chemistry. The Angiosperms353 v.1 target capture kit (Johnson et al., 2018) was purchased from Arbor Biosciences (Arbor Biosciences, Ann Arbor, Michigan, USA) and used for targeted enrichment of each sample. Two 8-reactions of the Angiosperms353 probes were used, which resulted in six libraries for each sample.

### 2.3 Sequence cleaning, assembly, and alignment

All analyses were run on the HiPerGator SLURM supercomputing cluster housed at the University of Florida, Gainesville, Florida, USA. Raw sequences were filtered and had adapters trimmed using SECAPR v1.1.12 (Andermann et al., 2018; Faircloth, 2016). After adjusting parameters and re-running SECAPR several times, as performed in the tutorial (https://htmlpreview.github.io/?https://raw.githubusercontent.com/AntonelliLab/seqcap_processor/master/docs/documentation/subdocs/cleaning_trimming.html), we required a minimum length of 60, a simple clip threshold of 4, a palindrome clip threshold of 20, a trailing quality of 40, seed mismatches 4, and a head crop of 10. Reads were assembled using HybPiper v1.3.1 (Johnson et al., 2016) with default settings, except BWA (Li and Durbin, 2009) was used for read mapping. Samples with paralogs were investigated using the HybPiper scripts paraloginvestigator.py and paralogretriever.py. For each locus with paralog issues, we generated alignments using MAFFT v7.407 (Katoh and Standley, 2013) followed by tree reconstruction using FastTree v2.1.7 (Price et al., 2010). This resulted in 10 loci being removed from further processing. Supercontig sequences were assembled using the intronerate.py script in HybPiper. Sequence alignment of each individual supercontig was completed in MAFFT v7.407 (Katoh and Standley, 2013) using --auto, --ep 0.123, and --op 2. Each alignment had columns containing less than 15% occupancy removed using the pxclsq command of phyx (Brown et al., 2017). Alignment statistics were summarized using the summary command implemented in AMAS (Borowiec, 2016).

### 2.4 Phylogenetic Analyses

We constructed two datasets composed of supercontig sequences (i.e., consensus sequences for each targeted locus including intronic and exonic regions). The full dataset included all samples regardless of missing data. The second dataset included taxa with less than 70% missing data in the supermatrix alignment (i.e., <= 70% gaps/ambiguities; referred to as Taxa70), a threshold that has previously been shown to improve resolution (Shah et al., 2021). Both datasets were analyzed using a concatenated alignment with a corresponding partition file in IQ-TREE v2.0-rc1 (Nguyen et al., 2015), which were produced using pxcat from phyx (Brown et al., 2017). A partitioned phylogenetic analysis was performed using the GENESITE partitioning analyses (--sampling GENESITE) (Gadagkar et al., 2005; Hoang et al., 2018), which helps reduce chances of overestimating bootstrap support. We implemented the ultrafast bootstrap approximation (UFBoot2) (Hoang et al., 2018) as well as an SH-like approximate likelihood ratio test (SH-aLRT; (Guindon et al., 2010), each with 1000 replicates (-B 1000; -alrt 1000) to assess support. The best-fit partitioning scheme using the greedy algorithm of PartitionFinder (Lanfear et al., 2012) with a relaxed clustering percentage of 10 (Lanfear et al., 2014) followed by tree reconstruction (-m TESTMERGE; -rcluster 10) was used.

### 2.5 Discordance and concordance analyses

Recent studies have shown that high bootstrap support can be recovered for many clades despite a low number of genes supporting a topology (Minh et al., 2020; Pease et al., 2018). Therefore, we quantified discordance and concordance within our supercontig dataset using multiple measures. First, we examined gene concordance factors (gCF; percentage of genes supporting the input topology) and site concordance factors (sCF; percentage of sites informative for a branch) as implemented in IQ-Tree v2.0-rc1 (Minh et al., 2020). Gene trees were generated from the partition file (–S option) with 1000 ultrafast bootstraps to obtain support values across each gene tree. The resulting concatenated species trees from the maximum likelihood analysis in addition to the individual gene trees were used to calculate both gCF and sCF with 100 random quartets in the sCF analysis (--scf 100).

Quartet Sampling (QS) provides unique and more revealing information pertaining to the potential causes behind discordance often prevalent in phylogenomic datasets (Pease et al., 2018). QS measures quartet concordance (QC), quartet differential (QD), quartet informativeness (QI) and taxon concordance (quartet fidelity; QF) (Pease et al., 2018). Quartet concordance (QC) measures the relative support for a clade when comparing across quartets and can provide evidence for the existence of an alternative topology. Positive values indicate concordance with the focal tree and the sampled quartets, negative values indicate that a discordant topology is most favored, and values of 0 indicate equal support among the different topologies sampled. Quartet differential (QD) measures the amount of support for an alternative evolutionary history. QD values of 1 indicate no skew in the proportion (no alternative evolutionary history), and values of 0 indicate that all trees sampled come from one of the two alternative relationships. Quartet informativeness (QI) indicates the amount of informative information available for a branch. QI values of 1 indicate all quartets were informative, whereas values of 0 indicate low informativeness for the branch. Lastly, quartet fidelity (QF) is a different measure of taxon’s “rogue-ness” in a data set. We examined Quartet Sampling (QS) values for both the full dataset and Taxa70 dataset using the resulting phylogenies from IQ-Tree along with their corresponding alignments and partition files. We ran 300 repetitions with a log-likelihood difference cutoff of 2 using the RAxML-NG v0.9.0 engine (Kozlov et al., 2019).

We further quantified and visualized discordance using DiscoVista (Sayyari et al., 2018), which provides a number of visualizations for gene tree discordance. We specifically investigated the number of gene trees that supported various clades within the Ledebouriinae. We defined the four major clades recovered from IQ-Tree (i.e., Ledebouria Clade A, *Ledebouria* Clade B, *Resnova*, and *Drimiopsis*) and investigated the number of gene trees that supported their monophyly. We also defined different combinations of the four major clades and investigated the number of gene trees that supported their monophyly (e.g., *Ledebouria* + *Resnova, Ledebouria* + *Drimiopsis, Ledebouria* Clade A + *Ledebouria* Clade B, etc.). We also investigated the number of gene trees that supported the inclusion of two taxa/samples within different clades of the phylogeny. These taxa, which were *Drimiopsis botryoides* subsp. *botryoides* CCH153 Tanzania, and *Ledebouria* sp. 1 Mozambique (Fig. 4), were studied in detail based on the results from SVDQuartets (below). We used the Taxa70 gene trees as input for the DiscoVista visualizations using a bootstrap cutoff value of 85 but recognizing that UFBoot recommends values above 95 be considered as strongly supported.

### 2.6 Species tree estimation

To obtain a species tree while also accounting for potential instances of incomplete lineage sorting, we used ASTRAL III v5.6.2 (Zhang et al., 2018). We used the fully resolved gene trees for both the full and Taxa70 datasets that were built using maximum likelihood as implemented in IQ-Tree v2.0-rc1.

ASTRAL uses gene trees as input, which can compound errors made during the alignment and tree building steps and does not incorporate information found in the alignment itself. To address these concerns, we used singular value decomposition quartet species-tree estimation (SVDQuartets) as implemented in PAUP* v4.0a (Swofford, 2001), which directly uses the alignment files for species tree reconstruction under the multi-species coalescent model while accounting for processes such as incomplete lineage sorting (Chifman and Kubatko, 2014). We input both the full dataset and Taxa70 dataset, each with their corresponding partition files. We ran an exhaustive search and assessed support using 100 bootstrap replicates in PAUP* (Swofford, 2001).

### 2.7 Hybridization analyses

Based on a comparison of results from the species-tree analyses as well as manual gene tree inspection, we suspected that historical gene flow may have occurred within the Ledebouriinae (i.e., we found that *Drimiopsis botryoides* subsp. *botryoides* CCH153 was nested within *Ledebouria* Clade A in 36% of the gene trees). To investigate potential reticulation within this clade, we used a pseudolikelihood approach as implemented in SNaQ (Solís-Lemus and Ané, 2016). This method estimates a network while accounting for incomplete lineage sorting and allowing for gene flow to occur. We ran three separate analyses allowing for zero to three hybridization events (hmax = 0, 1, 2). We used the log pseudolikelihood profile of these runs to distinguish between the best fitting model. As input, we used the Taxa70 gene trees from IQ-Tree to limit noise introduced by missing data. Hybridization analyses are still computationally demanding on large datasets (20+ tips), therefore, we subsampled ten tips from the phylogeny and conducted ten separate analyses on these reduced datasets. We included the suspected *Drimiopsis* CCH153 sample in each analysis as well as one additional randomly selected tip from the *Drimiopsis, Resnova*, and *Ledebouria* Clade B, and one outgroup taxon (i.e., *Massonia*). Because the suspected *Drimiopsis* sample was frequently nested within *Ledebouria* Clade A, each of our replicates included five random samples from this clade as a way to more precisely determine the location within *Ledebouria* Clade A where the hybrid edge(s) may occur. All replicate SNaQ analyses were conducted with 10 replicates on a random starting tree. Quartet concordance factors were summarized from gene trees inferred with IQ-Tree.

## 3. Results

### 3.1 Sequence capture

Of our 96 samples submitted for sequencing, 95 were returned. One herbarium specimen failed library preparation. An additional three samples failed to return sufficient coverage. Of the remaining 92 samples, percent on-target reads ranged from 1.9% to 29% (mean = 11%). The number of genes with contigs ranged from 12 – 348 (mean = 298). The number of genes per sample with sequences ranged from 4 – 348 (mean = 254). From the 353 targeted genes, we recovered 335 for our 92 samples after removing loci with insufficient data and paralogy concerns. Sample completeness per locus ranged from three to 91 samples with no gene recovered for every sample. Average supercontig length was 739 bp (min = 96, max = 2796). After alignment and trimming, individual gene alignment length ranged from 339 to 7593 bp, missing data per alignment was 24 – 63%, proportion of variable sites ranged from 6.4 – 95.3%, and proportion of parsimony informative sites ranged from 0 – 82.2%. Overall, herbarium samples contained the highest percentage of missing data compared to samples obtained from silica-dried leaf tissue (Fig. S1; Table S2).

### 3.2 Maximum likelihood and species tree estimation

Regardless of the full dataset including a significant amount of missing data, the four major clades (i.e., *Ledebouria* Clade A, *Resnova, Ledebouria* Clade B, *Drimiopsis*) received overall high support according to SH-aLRT, ultrafast bootstrap (UFBoot), and local posterior probability (LPP) (Table 1, Fig. 3, Figs. S1 – S4). Our analyses consistently recovered a polyphyletic *Ledebouria*, with each *Ledebouria* clade sister to either *Resnova* or *Drimiopsis* (Fig. 2). However, two results warrant attention. First, in the full dataset, we found a polyphyletic *Resnova* (Fig. S1) with low LPP (Fig. S3). After the removal of one herbarium sample, *Resnova lachenalioides*, that consistently jumped across clades potentially due to missing data (compare Figs. S2, S3, S10), we found increased support for a monophyletic *Resnova* across all estimates (Table 1, Fig. 3, Figs. S4, S5). Second, *Drimiopsis* + *Ledebouria* Clade B recovered overall high support except in the full dataset (Table 1, Figs. S2, S3). Using all samples, we found low support for *Ledebouria* Clade B in terms of the placement of *Ledebouria* sp. 1 Mozambique (Fig. 3, Table 1, Figs. S2, S3). The subclade sister to this sample shows higher but still low support in the full dataset (Figs. S1, S2). In the Taxa70 dataset, we find stronger support for *Drimiopsis* + *Ledebouria* Clade B (Figs. S3, S4). Support increases further for *Drimiopsis* + *Ledebouria* Clade B, as well as for *Ledebouria* Clade B after the removal of *Ledebouria* sp. 1 Mozambique (Table 1, Figs. S6, S7). Some of the lowest SH-aLRT and UFBoot support values are found along the backbone of a subclade in *Ledebouria* Clade A (i.e., the clade sister to *Ledebouria* sp. CCH003). Similar results are also shown in the ASTRAL topology with some of the lowest LPP values, the same shortest internal branches, and differences in topology compared to the maximum likelihood tree (Fig. 3; Figs. S3, S5).

**Figure 3.**
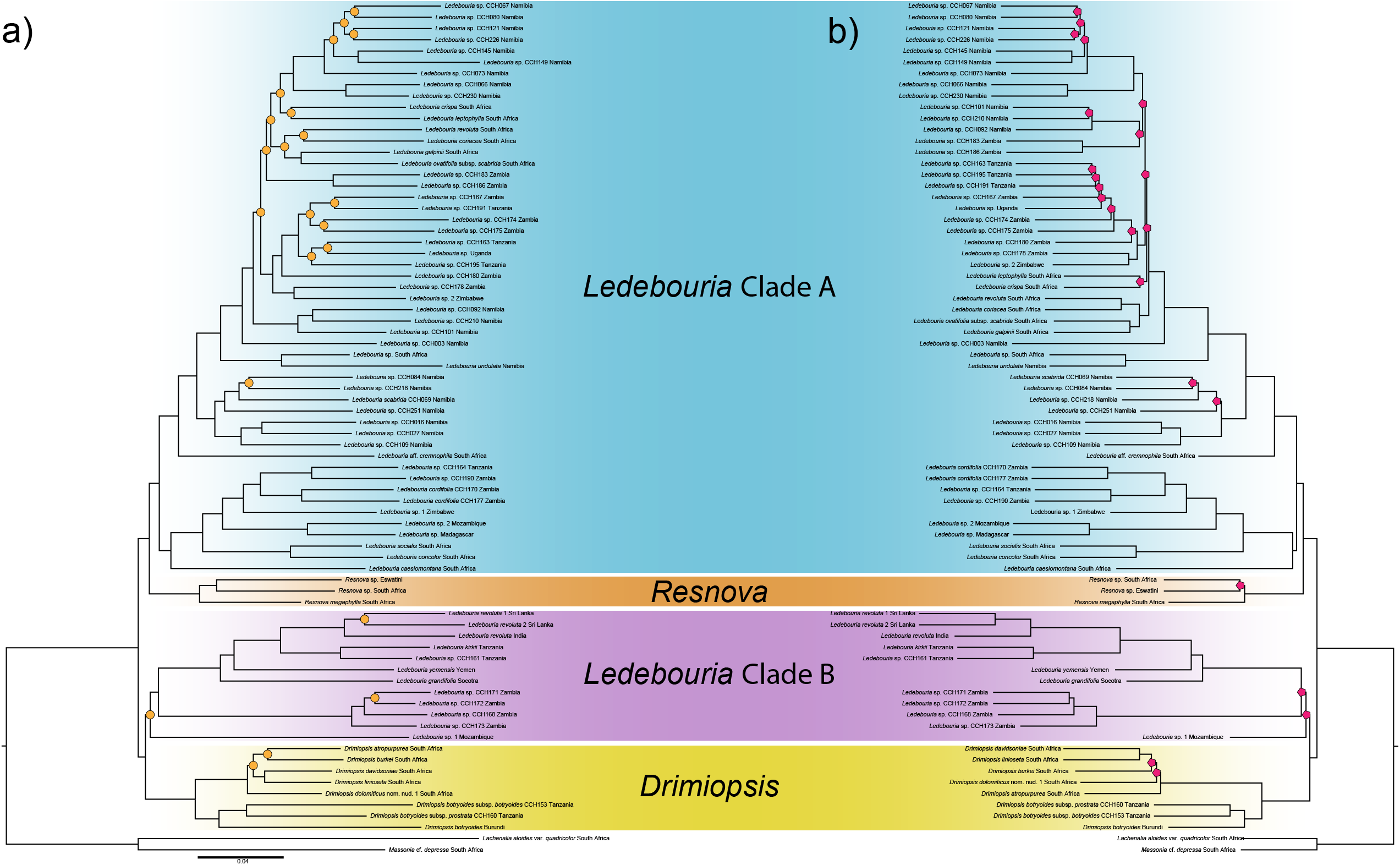
Phylogenetic reconstruction of the Taxa70 datasets using maximum likelihood as implemented in IQ-Tree (a), and species tree estimation as implemented in ASTRAL III (b). a) Orange circles indicate nodes with SH-aLRT and ultrafast bootstrap values below 80 and 95, respectively. b) Pink polygons indicate nodes with local posterior probability (LPP) below .95. Note, the same four major clades are recovered in both, with differences in relationships of subclades and tips between the two analyses, particularly within *Ledebouria* Clade A.

**Table 1.**
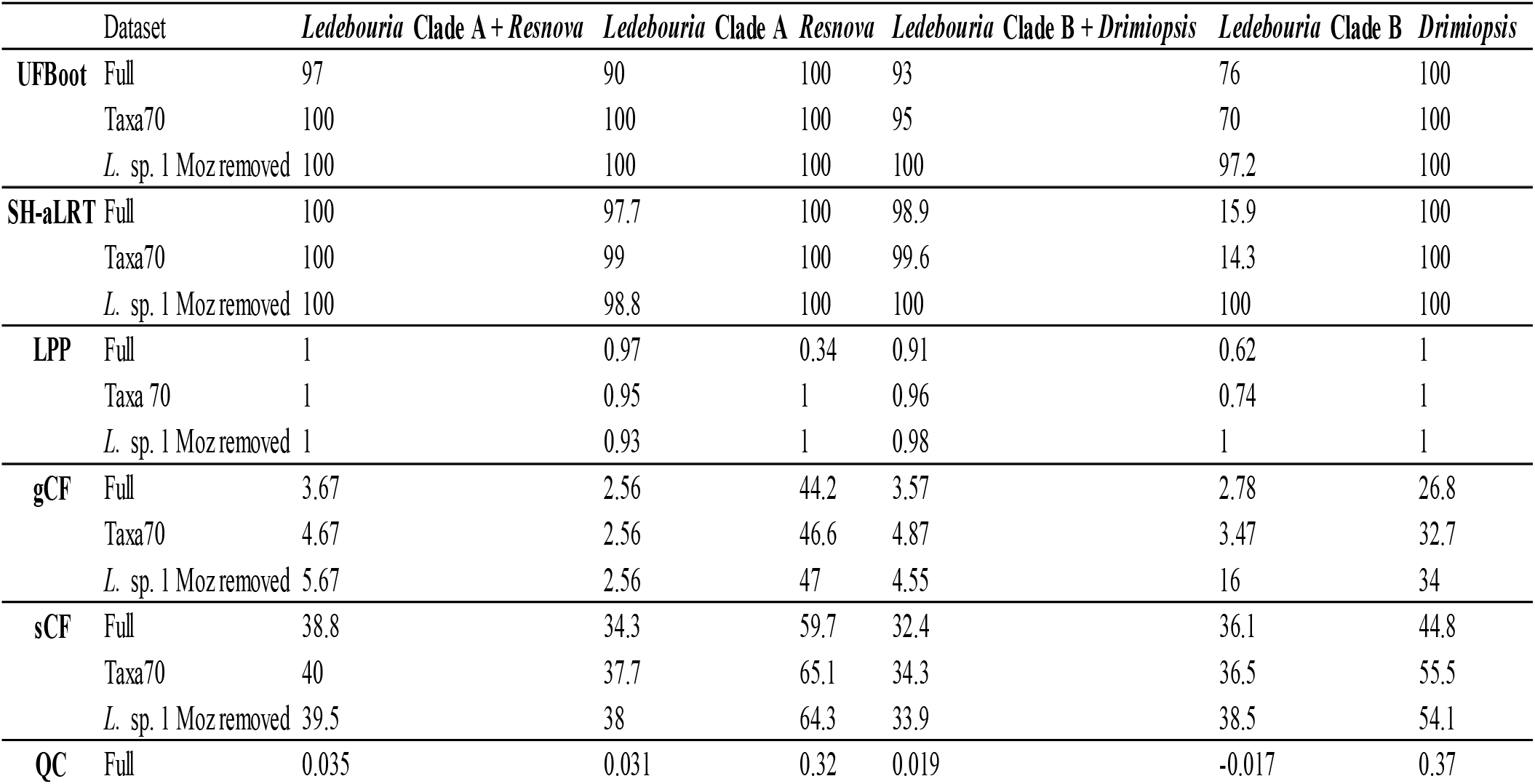
Support value measures, gene concordance factor (gCF), site concordance factor (sCF), and quartet sampling measures for the four major clades recovered. Full shows values for the full dataset, Taxa70 reports values for the Taxa70 dataset, and *L*. sp. 1 Moz removed shows values when *Ledebouria* sp. 1 Mozambique is removed from the analyses. Support measures are ultrafast bootstrap (UFBoot), SH-like approximate likelihood ratio test (SH-aLRT), and local posterior probability (LPP). Support measures are those recovered using the GENESITE partition resampling measure in IQ-Tree. Gene concordance factor (gCF) is expressed as the percentage of genes that support a given topology, site concordance factor (sCF) is expressed as the number of sites informative for a branch. Quartet sampling measures include quartet concordance (QC), quartet differential (QD), and quartet informativeness (QI). See Pease et al. (2018) for a detailed explanation of these measures.

SVDQuartets largely agrees with IQ-Tree and ASTRAL for both the full dataset and Taxa70 dataset (Figs. S10, S12). However, *Resnova lachenalioides* was nested within *Ledebouria* Clade B in the full dataset (Fig. S10). In both the full dataset and Taxa70 dataset, *Ledebouria* sp. 1 Mozambique was placed sister to the remaining Ledebouriinae. Topological differences were found between the SVDQuartet tree and the SVDQuartet bootstrap consensus tree in the Taxa70 dataset (Figs. S12, S13). The Taxa70 SVDQuartet bootstrap tree recovered *Drimiopsis* as sister to *Resnova* + *Ledebouria* Clade A, but with the lowest bootstrap support of all the nodes (Fig. S13).

### 3.3 Discordance and concordance

We found relatively low gene (gCF) and site (sCF) concordance factors for many nodes in both the full and Taxa70 datasets (Table 1; Figs. S1, S4). A notable improvement in support values was recovered for *Ledebouria* Clade B with the removal of *Ledebouria* sp. 1 Mozambique (Table 1; Fig. S6). Quartet concordance values were mostly positive for the four major clades (Table 1; Figs. S8, S9), but values were low for most backbone branches indicating close to equal levels of concordance and discordance among the quartets for the branch of interest Quartet differential (QD) values were regularly above .5, indicating somewhat similar frequencies of the three possible topologies for a branch (Table 1; Figs. S8, S9). Quartet informativeness (QI) indicated low informativeness of the quartets for the topology, reflecting the results of sCF (Table 1; Figs. S8, S9). DiscoVista reported only one gene tree that supports a monophyletic *Ledebouria* (Fig. 4b; Table S3). 126 gene trees supported a monophyletic *Resnova*, 103 genes supported a monophyletic *Drimiopsis*, nine genes supported *Ledebouria* Clade A, 16 genes supported a monophyletic *Ledebouria* Clade B. When *Ledebouria* sp. 1 Mozambique was excluded from *Ledebouria* Clade B, 47 genes supported its monophyly. 27 genes support *Ledebouria* sp. 1 Mozambique as an outgroup taxon. 17 gene trees support *Drimiopsis* + *Ledebouria* Clade B, and 16 gene trees support *Resnova* + *Ledebouria* Clade A. Full outputs plus clade definitions used as input for DiscoVista can be found in the Dryad repository (**FOR REVIEW ONLY**: https://datadryad.org/stash/share/hZprQ-ukfTvIxPNT9XSCe3WfZ0rGexqaNf3POk2egsQ).

**Figure 4.**
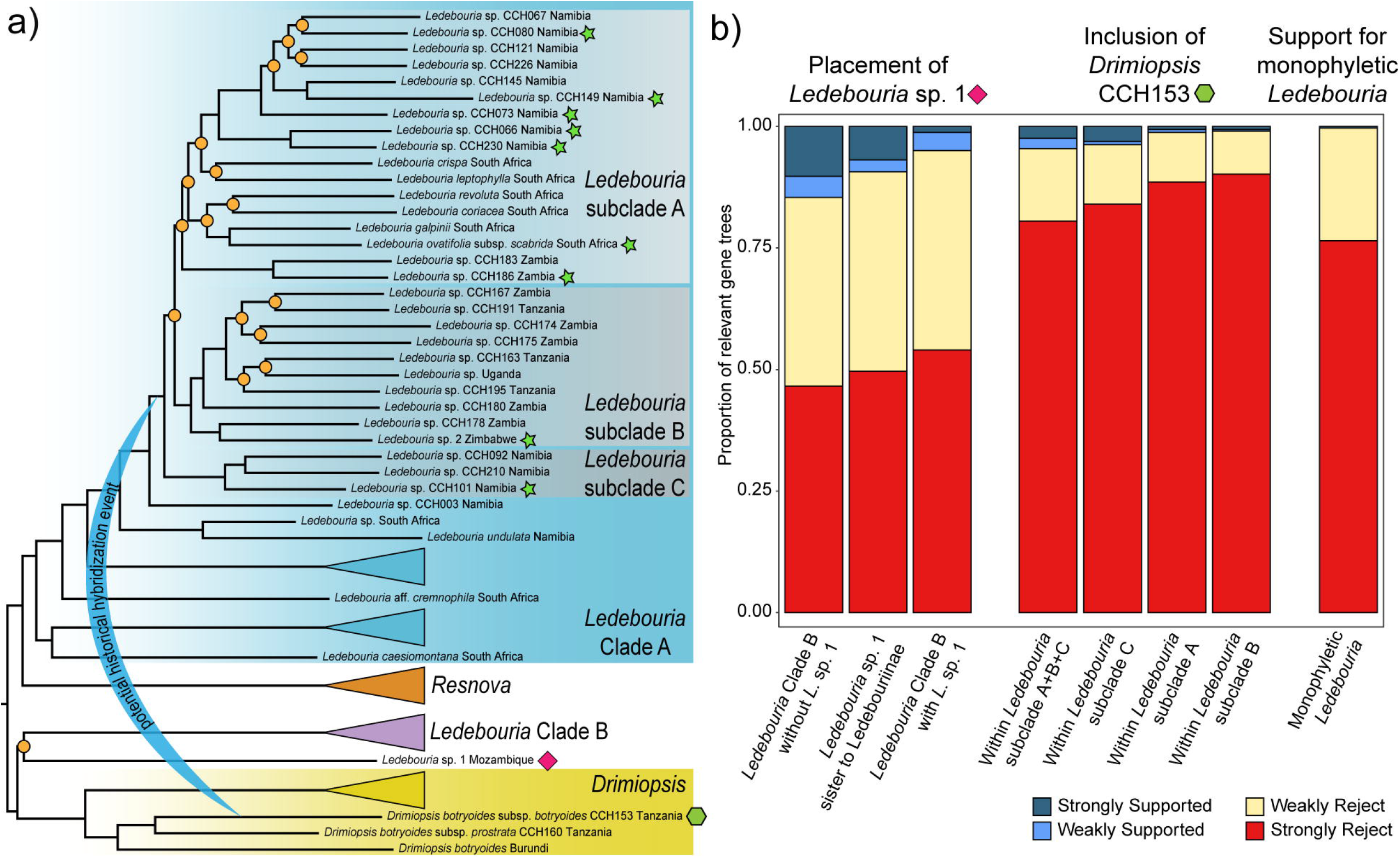
Summary of hypothesized hybridization and phylogenetic discordance. a) Maximum likelihood reconstruction with select clades collapsed for clarity; see Fig. 1 for the full Taxa70 phylogeny. Green stars indicate taxa identified as potentially involved in a hybridization event with *Drimiopsis botryoides* subsp. *botryoides* CCH153 [green polygon] in randomly subsampled replicate analyses. Given these results, we hypothesize an ancient hybridization event involving an ancestor of this subclade. Orange circles indicate nodes with SH-aLRT and ultrafast bootstrap values below 80 and 95, respectively. b) The number of relevant gene trees (i.e., those with the clade(s) of interest) that support or reject clades of interest. The first three bars show the number of gene trees that support the placement of *Ledebouria* sp. 1 Mozambique [phylogenetic placement marked by a pink diamond in a] in relation to *Ledebouria* Clade B and the Ledebouriinae. The middle four bars show the number of genes that support the placement of *Drimiopsis botryoides* subsp. *botryoides* CCH153 within the subclade of *Ledebouria* Clade A where an ancient hybridization event may have occurred. The last bar shows the number of gene trees that support a monophyletic *Ledebouria*.

### 3.4 Hybridization

We found support for one hybridization event in our dataset (Table S4). In all ten replicate analyses, a hybrid edge was inferred between *Drimiopsis botryoides* subsp. *botryoides* CCH153 and at least one taxon within *Ledebouria* Clade A. This compliments the DiscoVista results, which showed that a surprisingly high number of gene trees placed *Drimiopsis botryoides* subsp. *botryoides* CCH153 within a subclade of *Ledebouria* Clade A (Fig. 4b; Table S3). Each of the *Ledebouria* Clade A samples implicated in gene flow mostly fall within a single subclade (Fig. 4). Interestingly, this *Ledebouria* subclade also has the lowest supported nodes within the entire phylogeny (Fig. 3, 4a). If multiple samples from this subclade were included, often only one sample was inferred as being involved in hybridization. We suspect the high amount of discordance within this subclade to be contributing to the inability of the analysis to confidently pinpoint where the potential hybridization event occurred.

## 4. Discussion

We provide the first phylogenomic insights into the Ledebouriinae. Four major clades were recovered: *Ledebouria* Clade A, *Resnova, Drimiopsis*, and *Ledebouria* Clade B (Figs. 1b, 3, Figs. S2 – S7). The low number of informative genes coupled with coalescence and concordance analyses suggest that incomplete lineage sorting (likely due to rapid radiations) along the backbone of the group as indicated by short branch lengths, may have played a significant role in the group’s evolutionary history, a common theme in modern-day phylogenomic studies (Morales-Briones et al., 2021; Stubbs et al., 2020; Thomas et al., 2021). Our analyses also hint that historical hybridization may be contributing to further phylogenetic discordance within the group, particularly within a subclade of *Ledebouria* Clade A (Fig. 4).

Past phylogenetic studies of the Ledebouriinae have recovered conflicting signal or low resolution along the backbone of the group (Lebatha et al., 2006; Manning et al., 2003; Pfosser, 2012; Pfosser et al., 2003; Wetschnig et al., 2007). The focus of many of these studies has largely been on whether *Drimiopsis* and *Resnova* constitute two distinct taxa or are deeply nested within *Ledebouria* (Lebatha et al., 2006; Manning and Goldblatt, 2012; Manning et al., 2003). The cladistic analysis of Lebatha et al. (2006) using morphological and molecular data supported three distinct entities, and our results support the delimitation of *Drimiopsis* and *Resnova*. Both lineages have easily distinguishing characteristics, such as tepal shape, stamen positioning, and non-stipitate ovaries (Lebatha et al., 2006; Lebatha, 2004). However, in Lebatha et al. (2006), *Ledebouria* sampling was restricted to South African taxa. When including additional *Ledebouria* samples, multiple studies also found support (although weak) for a monophyletic *Drimiopsis* and/or *Resnova*, but they also hinted at potential unknown lineage diversity within *Ledebouria* as indicated by multiple clusters of *Ledebouria* samples (Pfosser et al., 2003; Stedje, 1998; Wetschnig et al., 2007). For example, Wetschnig et al. (2007) recovered two *Ledebouria* clades, each one sister to either *Resnova* or *Drimiopsis*, with *Ledebouria floribunda* as sister to the remaining Ledebouriinae. Here, we recovered a surprisingly similar topology, but the *Ledebouria* clade containing samples from India and Madagascar are sister to *Drimiopsis*, not *Resnova* as recovered by Wetschnig et al. (2007) (Fig. 1). Our results, along with past analyses, suggest that perhaps a more appropriate goal from the start should have been uncovering diversity within *Ledebouria* rather than debating the status of *Drimiopsis* and *Resnova*. We recommend that attention should now shift to delimiting and interpreting the history of the independent *Ledebouria* lineages as well as uncovering and describing more diversity. Reconfiguring both the broad- and fine-scale taxonomy of the Ledebouriinae is beyond the scope of this work, but our results lead us to question the status of *Ledebouria* sensu Manning (2003).

The current geographic distribution of the Ledebouriinae is reflected within the phylogeny and provides clues as to potential historical factors that have shaped their evolution. For example, the two *Ledebouria* clades overlap geographically in eastern Africa, which serves as a melting pot, center of diversity, area of endemism, and biogeographical hub for many different African plants and animals (Dagallier et al., 2020; Lebatha, 2004; Lorenzen et al., 2012; Zhou et al., 2011). Similar processes that have potentially shaped the historical evolution of the two *Ledebouria* clades (e.g., mountain building/increased aridity in eastern Africa) may have also shaped the current diversity and distribution of *Drimiopsis*, which, based on our limited sampling, consist of two clades: a South African clade, and a northern African clade (Fig. 1). These two *Drimiopsis* clades can be distinguished by morphological traits (i.e., tepal morphology) and ploidy level (i.e., East African *Drimiopsis* are polyploids) (Lebatha, 2004; Stedje and Nordal, 1987). Surprisingly, SVDQuartets consistently, albeit with low support, placed East African *Drimiopsis botryoides* subsp. *botryoides* (CCH153) within *Ledebouria* Clade A. Our network analysis implicates this same taxon in a potential hybridization event involving an ancestral taxon within *Ledebouria* Clade A (Fig. 4a; Fig. S13), and 15 gene trees support and 48 gene trees weakly reject the placement of this *Drimiopsis* within the *Ledebouria* subclade (Fig. 4b; Table S3). Today, sympatric populations of all Ledebouriinae can be found in sub-Saharan Africa, with the highest levels in eastern South Africa (Lebatha, 2004; Venter, 1993). Therefore, it is not unreasonable to hypothesize that ancestral populations of *Drimiopsis* and *Ledebouria* Clade A also overlapped in distribution leading to potential hybridization. The results of our preliminary hybridization analysis and the knowledge that East African *Drimiopsis* are known polyploids (Stedje and Nordal, 1987), leads us to suspect that hybridization may be at play in these lineages. However, much greater sampling of *Drimiopsis* is needed to fully test any hypotheses regarding historical hybridization within the Ledebouriinae, but our results certainly provide areas worthy of further investigation.

Ledebouriinae taxa are continually being described (Cumming, 2018; Hankey, 2020; Hankey et al., 2014), and enormous diversity remains to be introduced to the scientific community (Fig. 2). For example, *Ledebouria scabrida* Jessop is the only taxon currently described as endemic to Namibia (Jessop, 1973), but extensive fieldwork and our phylogenomic results show that far more species diversity awaits formal description (Howard, 2014). This is exemplified by *Ledebouria* sp. 1 Mozambique, whose placement has 1) low support in the IQ-Tree and ASTRAL trees (Fig. 3), 2) has different placement in the IQ-Tree and SVDQuartet trees (Fig. 3a, Figs. S10, S11), 3) has the lowest quartet fidelity value (i.e., a measure for rogue taxa) of all the tips (Taxa70 dataset; Fig. S15), 4) is found sister to the Ledebouriinae in several gene trees (Fig. 4b; Table S3), and 5) exhibits unique morphological characteristics compared to currently known Ledebouriinae (e.g., seeds possess a rostrate hilum giving them a distinctive pointed appearance; Fig. S16). Whether this accession represents a distinct Ledebouriinae lineage requires further sampling and morphological characterization and is just one example of the diversity awaiting description within the Ledebouriinae.

### 4.1 Conclusion

In this paper, we showcase the first phylogenomic analysis of the Ledebouriinae with the most comprehensive taxon sampling to date. Our results showcase the value of extending the phylogenomic repertoire even to groups with unexplored potential. Our use of the Angiosperms353 probe set was vital for uncovering a revealing Ledebouriinae phylogeny, but a custom probe set may prove even more informative. Our analyses suggest that gene tree conflict due to incomplete lineage sorting and hybridization is prevalent within the phylogeny (Table 1; Fig. 4), and several short backbone branches coupled with concordance factors suggest rapid radiations (Fig. 3). Assessing the individual contribution of incomplete lineage sorting and hybridization to the discordance found within the Ledebouriinae phylogeny remains to be fully addressed. Increasing *Ledebouria* samples from Central and West Africa as well as significantly improving sampling of *Resnova* and *Drimiopsis* will be necessary to fully capture the intricate genomic evolution of the group. The role that incomplete lineage sorting and hybridization have played in obscuring the evolutionary history of the Ledebouriinae present an exciting opportunity for further study.

## CRediT authorship contribution statement

**Cody Coyotee Howard**: Conceptualization, Methodology, Investigation, Resources, Data curation, Writing – Original Draft, Visualization, Funding acquisition. **Andrew T. Crowl**: Methodology, Software, Writing – Review & Editing, Visualization. **Timothy S. Harvey**: Conceptualization, Resources, Writing – Review & Editing. **Nico Cellinese**: Conceptualization, Writing – Review & Editing, Visualization, Supervision, Project administration, Funding acquisition.

## Declaration of interest

None

## Funding

This work was supported by funds from the Huntington Botanical Gardens; the Cactus and Succulent Society of America; the Pacific Bulb Society (Mary Sue Ittner Bulb Research Grant); the San Gabriel Cactus and Succulent Society; the Florida Museum of Natural History; the American Society of Plant Taxonomists; the University of Florida International Center; the University of Florida Department of Biology; the Botanical Society of America; the Society of Systematic Biologists; Xeric Growers; and numerous private donors.

## Acknowledgements

We are extremely grateful to Leevi Nanyeni, Silke Rügheimer and Dr. Esmeralda Klassen at the National Botanical Research Institute in Windhoek, Namibia for their assistance with fieldwork and specimen export while in Namibia; Inge Pehlemann for enjoyable Namibian excursions in search of *Ledebouria* and others plants; Dr. David Chuba at the University of Zambia with assistance in acquiring collection and export permits for Zambia; Dr. Neduvoto Mollel at the Tropical Pesticides Research Institute in Arusha, Tanzania for help in obtaining collection and export permits for Tanzania. Sincere thanks to the governments of Namibia, Zambia, and Tanzania for issuing collection and export permits. Collections from Namibia were made under permit numbers 1784/2013, 1908/2014, 2056/2016, and 2185/2016. Zambian collections were made under permit number TJ/DNPW/101/13/18. Tanzanian collections were made under permit number 2017-22-NA-2016-247. Living plants were imported under USDA permit numbers P37-09-00910, P37-16-00181, and P37-16-01462. We also express our deep gratitude to Dylan Hannon, Gottfriend Milkuhn, Tom Cole, and Tom McCoy for donating leaf material from geographically crucial taxa. Lastly, we also are grateful to Killian Fleurial, Taylor La Val, and Emily Sessa for assistance while in the field.

## Disclosure statement

The authors have no conflict of interests to disclose.

## Data availability statement

The raw genomic reads for each sample used in this study are available on the SRA (BioProject ID: PRJNA721471). Supplementary Tables and Figures as well as alignment files, individual gene trees, and output files for each analysis are available on Dryad (DOI: **FOR REVIEW ONLY** (https://datadryad.org/stash/share/hZprQ-ukfTvIxPNT9XSCe3WfZ0rGexqaNf3POk2egsQ).

## Supplementary Tables

Supplementary Table 1. List of specimens used in this study. Specimens include only those returned after adjusting for sequences coverage and removing sequences with paralogy warnings.

Supplementary Table 2. Gene recovery statistics per sample as output from HybPiper.

Supplementary Table 3. Number of gene trees that support or reject defined clades in DiscoVista. See Fig. 3 for clades/samples defined in the analysis.

Supplementary Table 4. SNaQ likelihood values for each of the ten randomly sampled subsets of the Ledebouriinae phylogeny when allowing 0 (H0) – 2 (H2) hybrid edges. Figure S14 shows the ten subset phylogenies.

## Supplementary Figures

Supplementary Figure 1. Heatmap visualization of gene recovery per sample output from HybPiper.

Supplementary Figure 2. Maximum likelihood phylogenetic reconstruction using IQ-Tree of the full dataset. Branch labels show SH-aLRT support value/Ultrafast bootstrap support value/gene concordance factor/site concordance factor. Strong support is branches with SH-aLRT and UFBoot values above 80 and 95, respectively.

Supplementary Figure 3. Species tree estimation of the full dataset inferred from ASTRAL. Branch labels indicate local posterior probability (LPP) support values.

Supplementary Figure 4. Maximum likelihood phylogenetic reconstruction using IQ-Tree of the Taxa70 dataset. Branch labels show SH-aLRT support value/Ultrafast bootstrap support value/gene concordance factor/site concordance factor. Strong support is branches with SH-aLRT and UFBoot values above 80 and 95, respectively.

Supplementary Figure 5. Species tree estimation of the Taxa70 dataset inferred from ASTRAL. Branch labels indicate local posterior probability (LPP) support values.

Supplementary Figure 6. Maximum likelihood phylogenetic reconstruction using IQ-Tree of the Taxa70 dataset with *Ledebouria* sp. 1 Mozambique removed from the analysis. Branch labels show SH-aLRT support value/Ultrafast bootstrap support value/gene concordance factor/site concordance factor. Strong support is branches with SH-aLRT and UFBoot values above 80 and 95, respectively.

Supplementary Figure 7. Species tree estimation of the Taxa70 dataset with *Ledebouria* sp. 1 Mozambique removed inferred from ASTRAL. Branch labels indicate local posterior probability (LPP) support values.

Supplementary Figure 8. Maximum likelihood phylogeny with quartet sampling values drawn on branches using the full dataset. Values on branches are quartet concordance (QC)/quartet differential (QD)/quartet informativeness (QI).

Supplementary Figure 9. Maximum likelihood phylogeny with quartet sampling values drawn on branches using the Taxa70 dataset. Values on branches are quartet concordance (QC)/quartet differential (QD)/quartet informativeness (QI).

Supplementary Figure 10. Species tree estimation of the full dataset using singular value decomposition quartet species-tree estimation (SVDQuartets). Phylogeny was reconstructed using an exhaustive search.

Supplementary Figure 11. Species tree estimation of the full dataset using singular value decomposition quartet species-tree estimation (SVDQuartets). Phylogeny was reconstructed using an exhaustive search. Branch labels indicate bootstrap support from 100 bootstrap replicates.

Supplementary Figure 12. Species tree estimation of the Taxa70 dataset using singular value decomposition quartet species-tree estimation (SVDQuartets).

Supplementary Figure 13. Species tree estimation of the Taxa70 dataset using singular value decomposition quartet species-tree estimation (SVDQuartets). Phylogeny was reconstructed using an exhaustive search. Branch labels indicate bootstrap support from 100 bootstrap replicates.

Supplementary Figure 14. SNaQ results from ten randomly sampled subsets of the Ledebouriinae phylogeny. Network 1 reports results from analyses that allowed 1 hybrid edge (h=1), and network 2 reports results from analyses that allowed up to 2 hybrid edges (h=2) for each replicate. Dark blue lines indicate the major hybrid edge, light blue lines indicate the minor edge. Inheritance probabilities are provided on the edges. Gray lines connect branches that are rotated across the two phylogenies.

Supplementary Figure 15. Maximum likelihood phylogeny showing quartet fidelity (QF) values for each of the samples in the Taxa70 dataset. Values represent a measure of taxon “rogueness”, that is, a measure of how much a tip jumps around a phylogeny.

Supplementary Figure 16. *Ledebouria* sp. 1 Mozambique (a) showing the pronounced rostrate hilum that sets it apart from other Ledebouriinae (e.g., image b) that typically have more spherical seeds.

## Notes

### Competing Interest Statement

The authors have declared no competing interest.

### Summary of Updates

Addressing reviewer's comments and edits; new title; new figure

