## Supplementary figures and images for "Peeling Back the Layers: First Phylogenomic Insights into the Ledebouriinae (Scilloideae, Asparagaceae)"

### Fig. S1

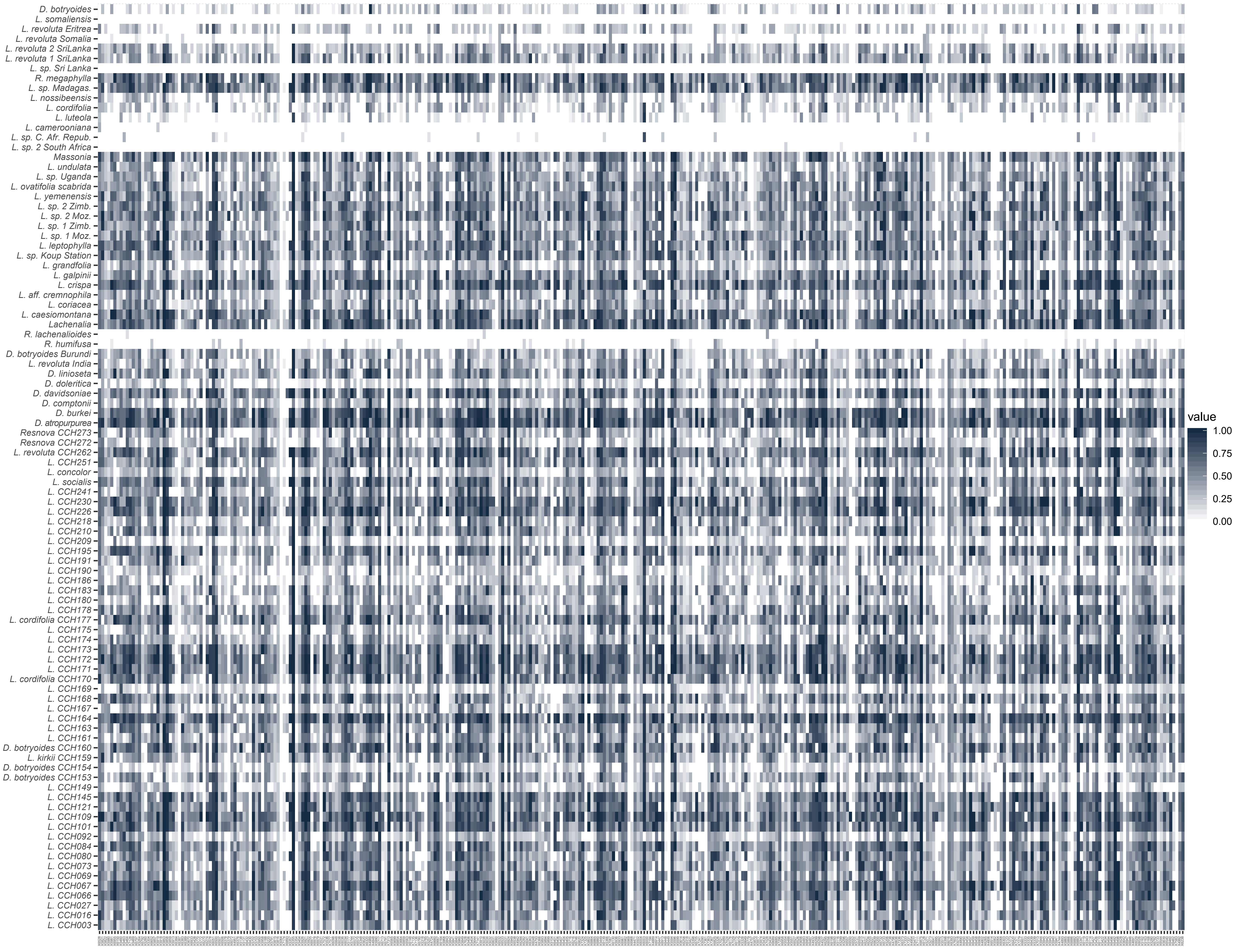

### Fig. S2

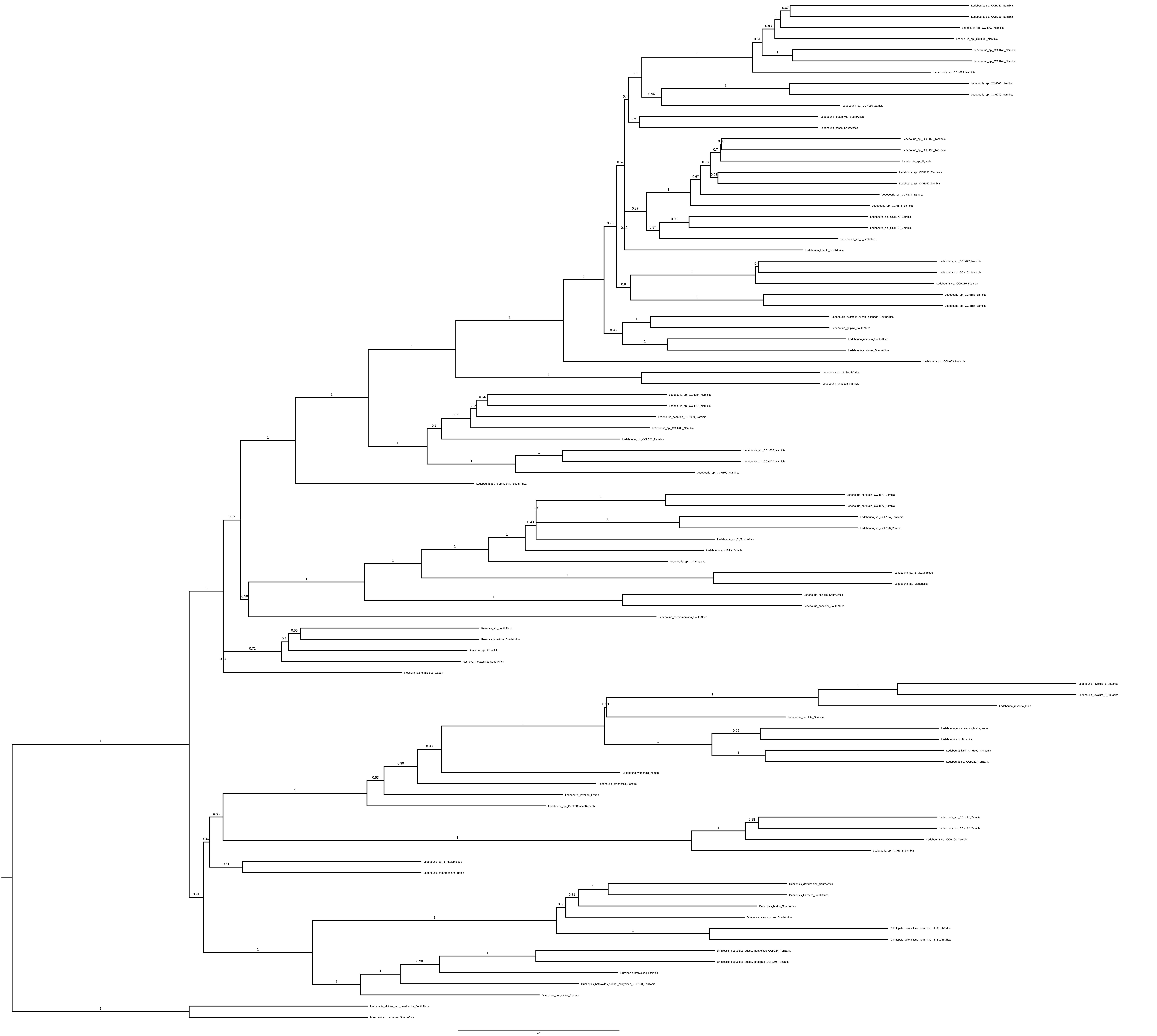

### Fig. S3

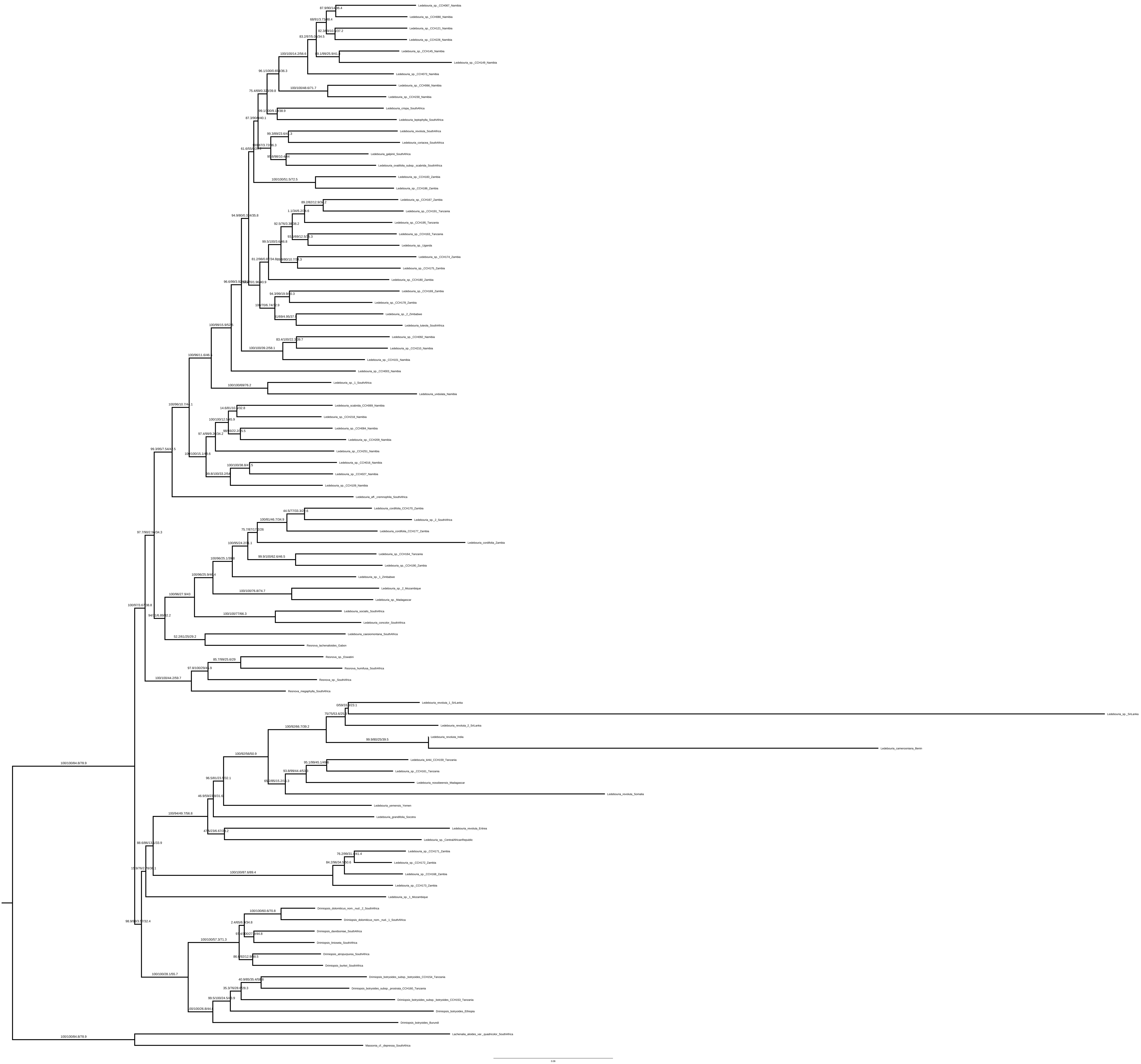

### Fig. S4

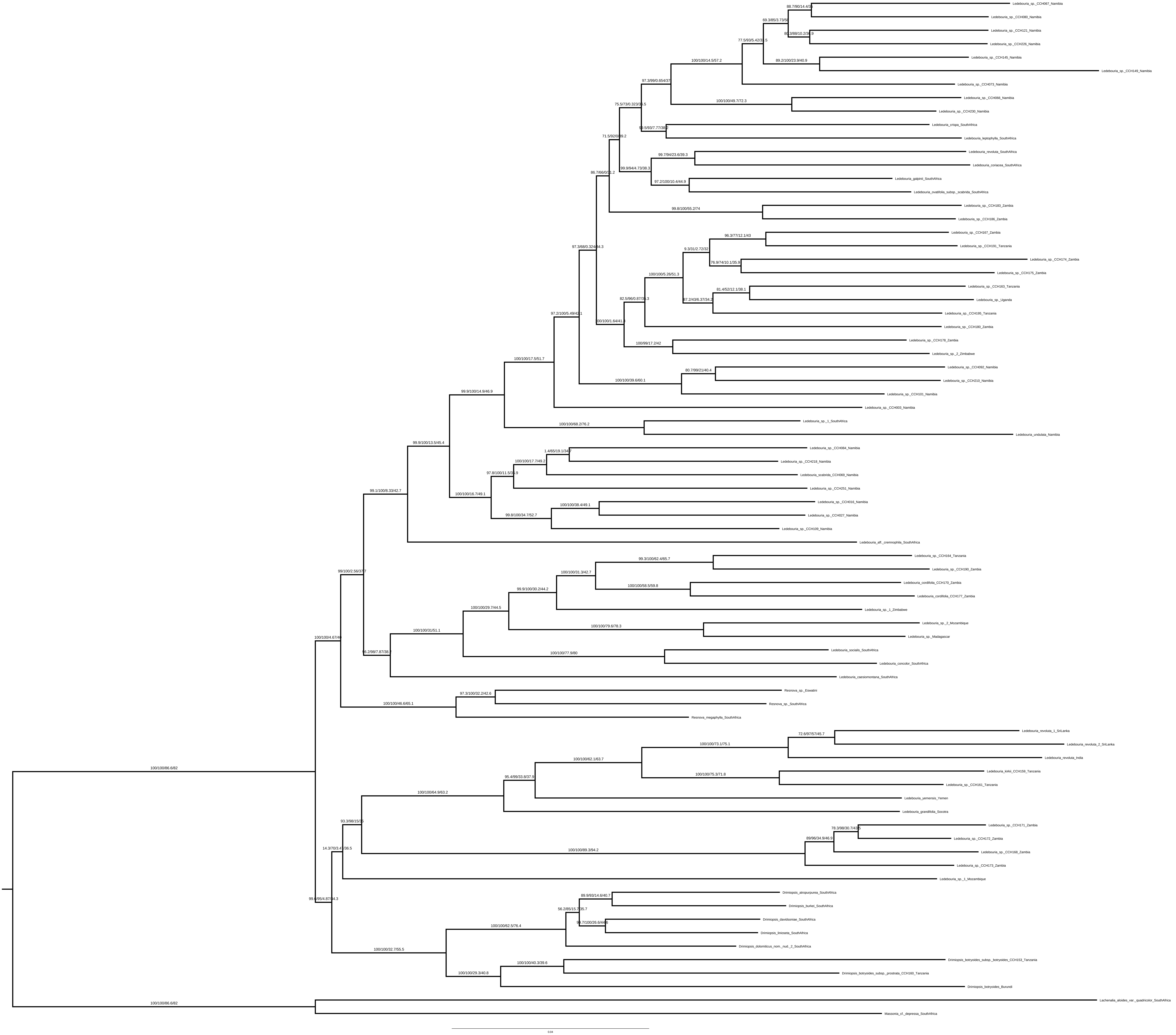

### Fig. S5

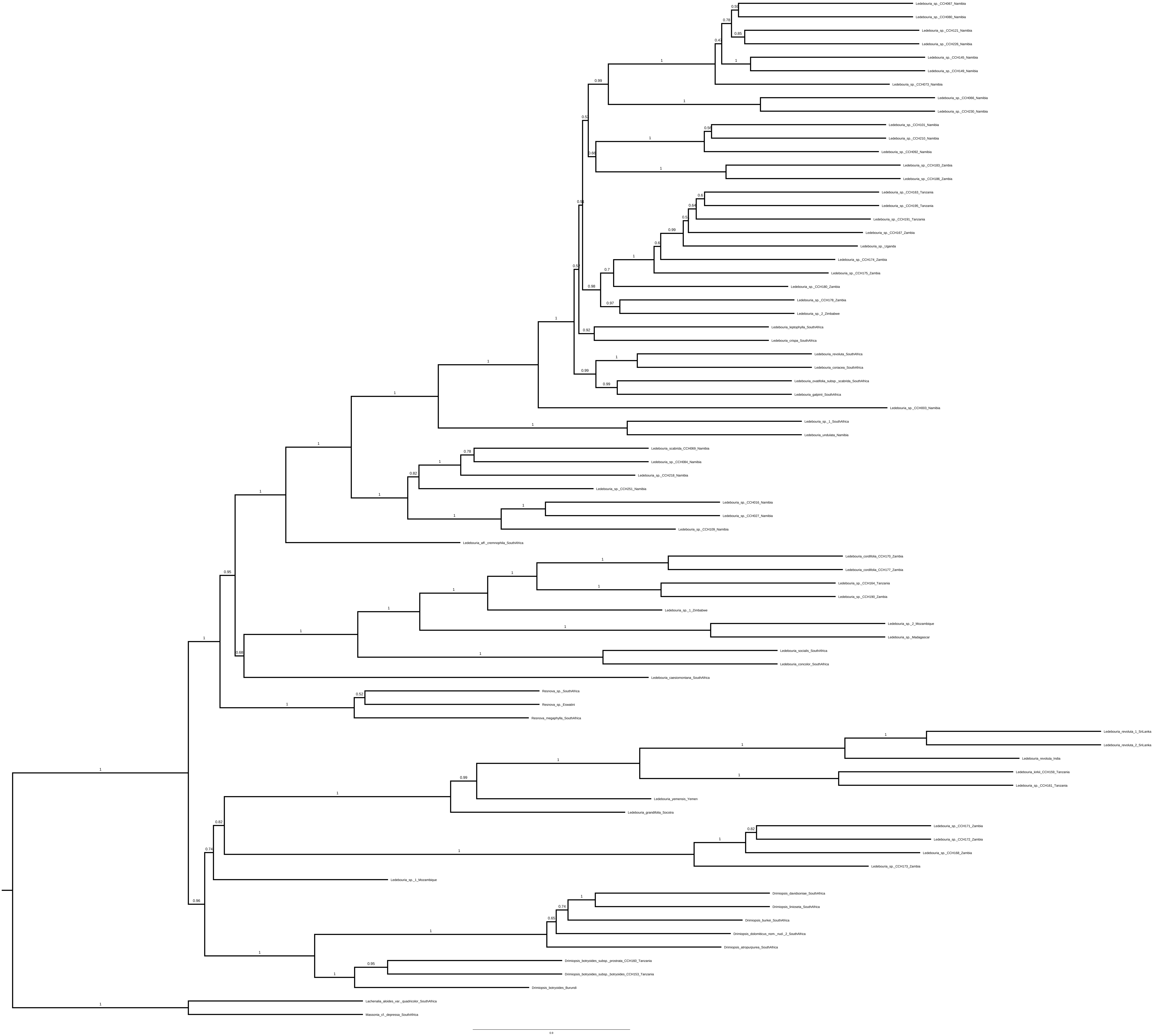

### Fig. S6

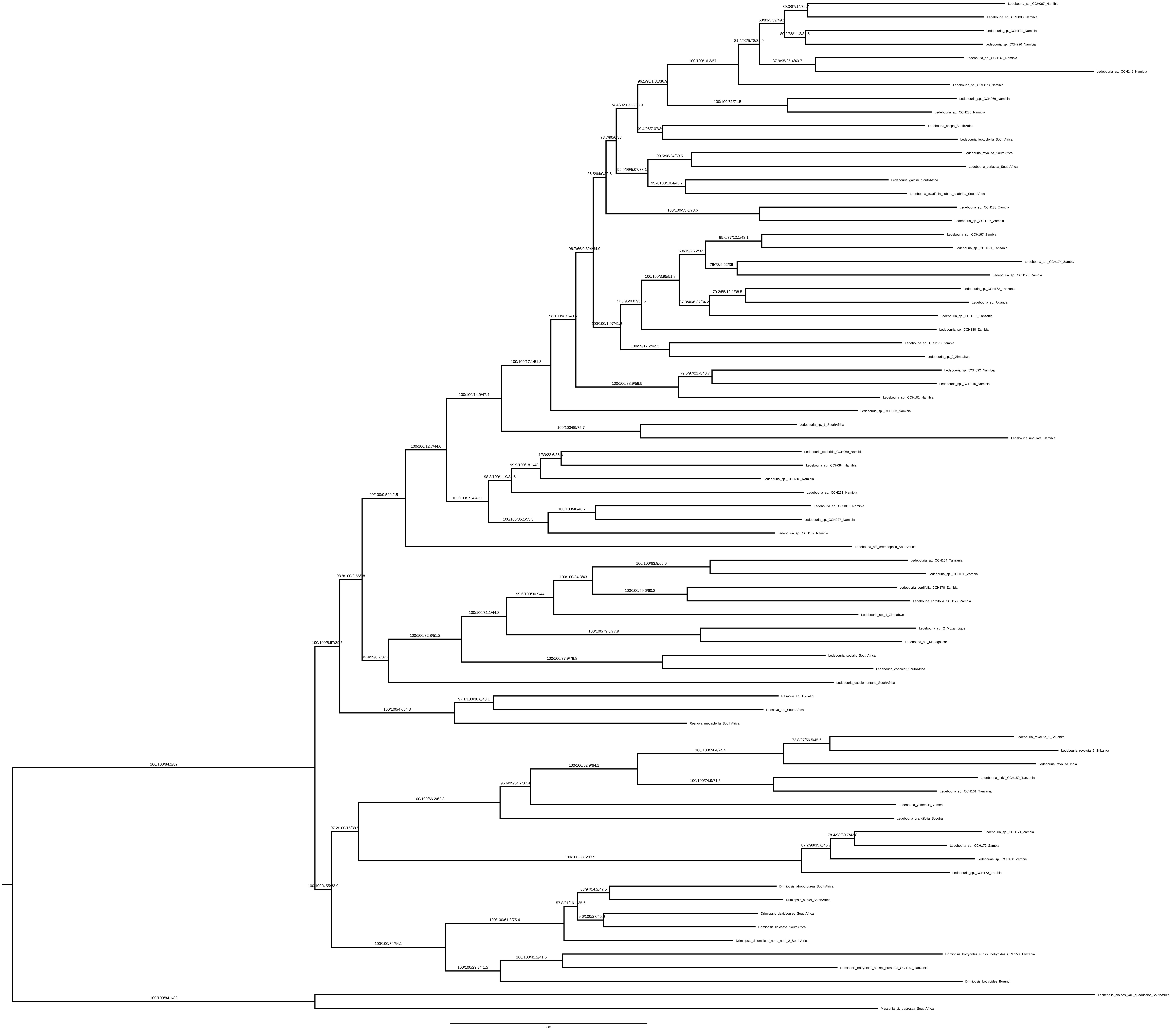

### Fig. S7

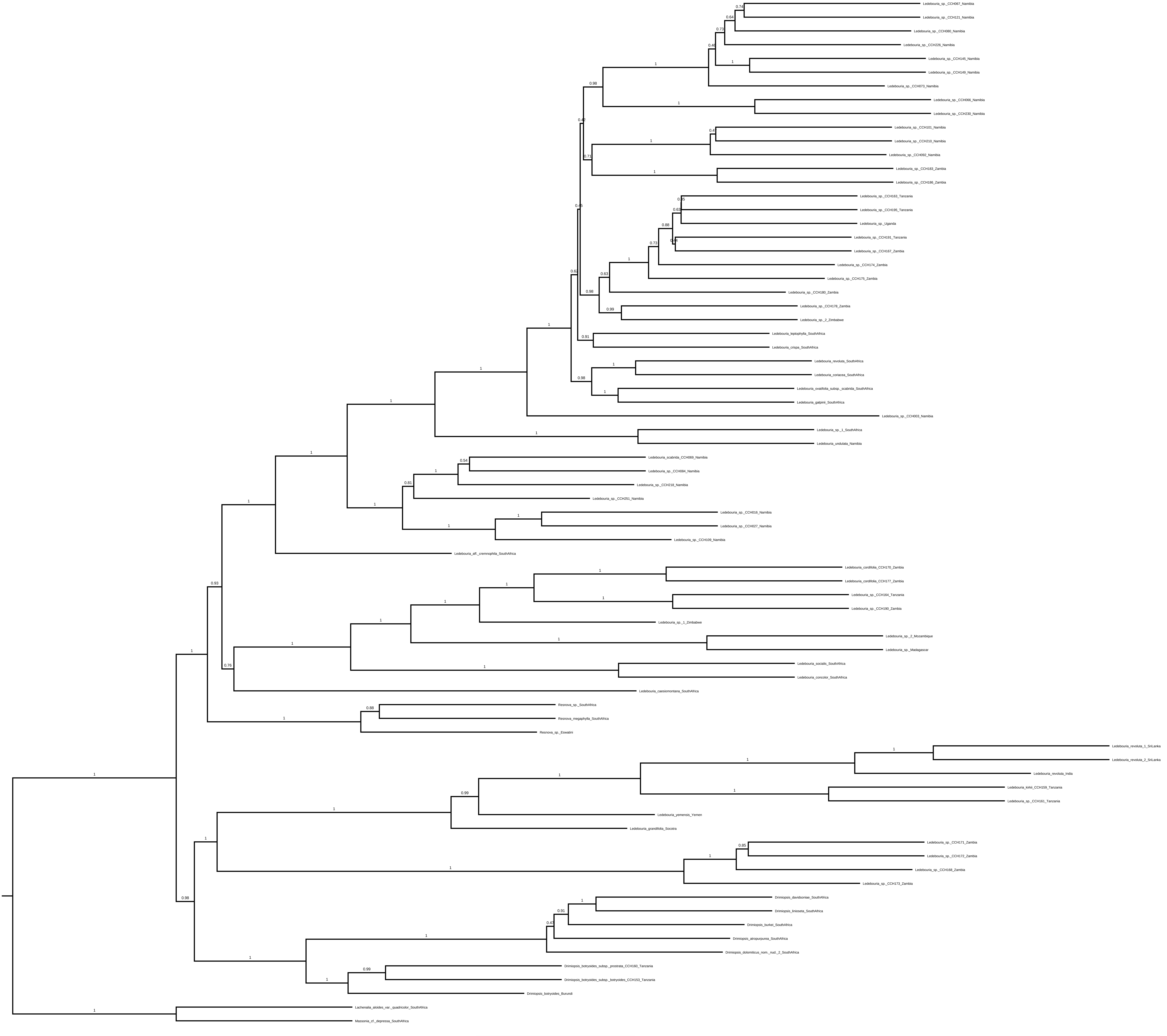

### Fig. S8

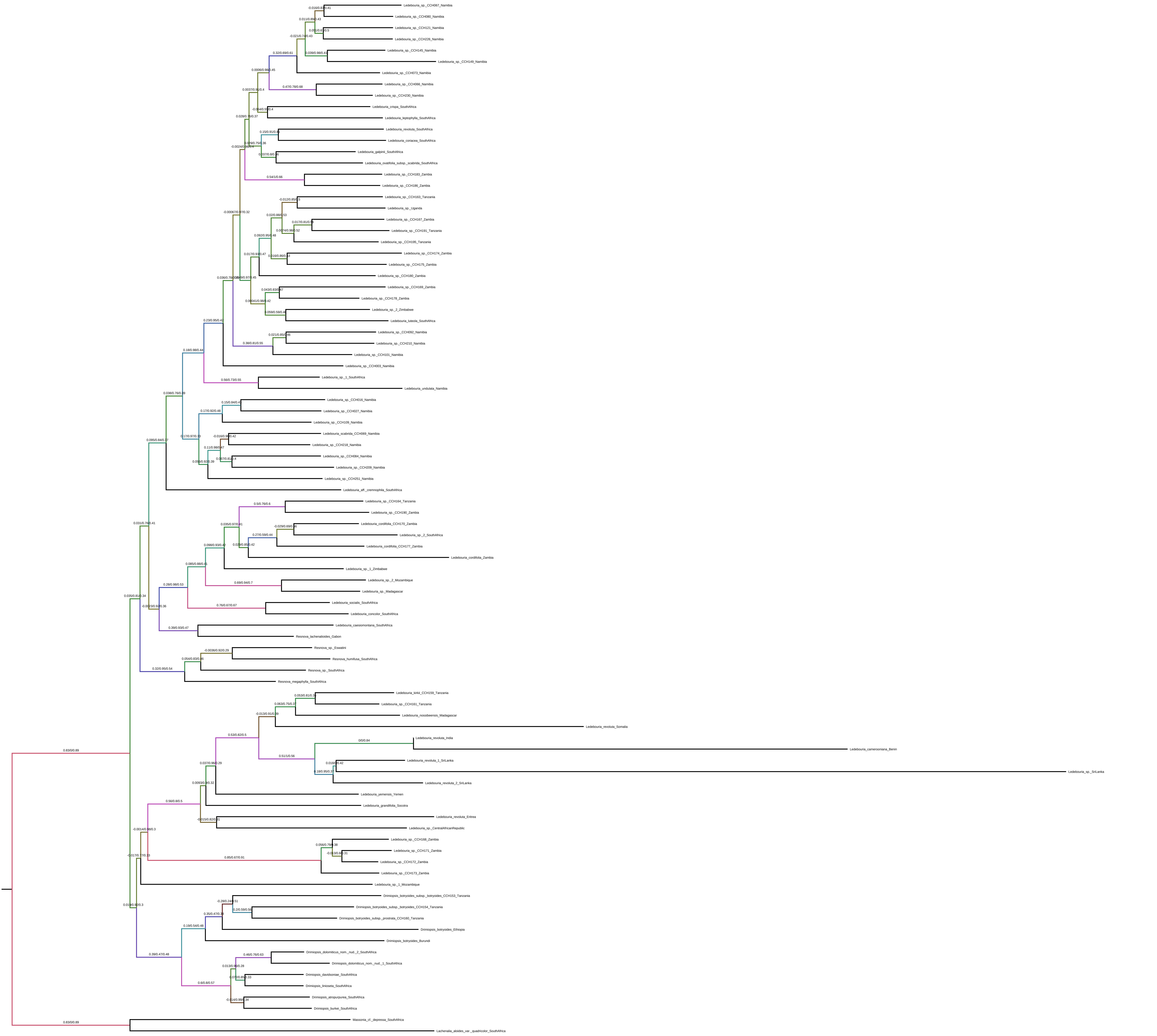

### Fig. S9

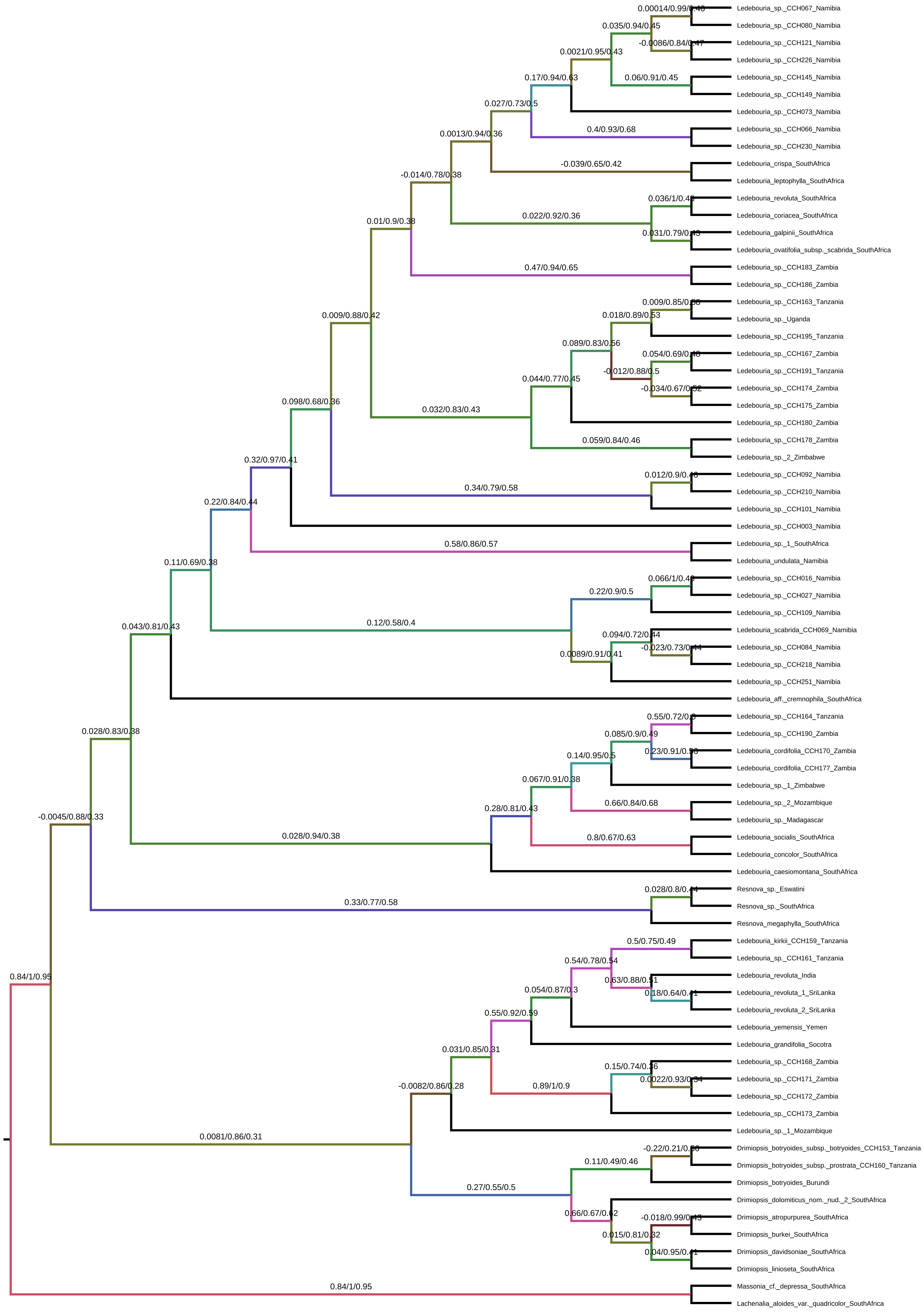

### Fig. S10

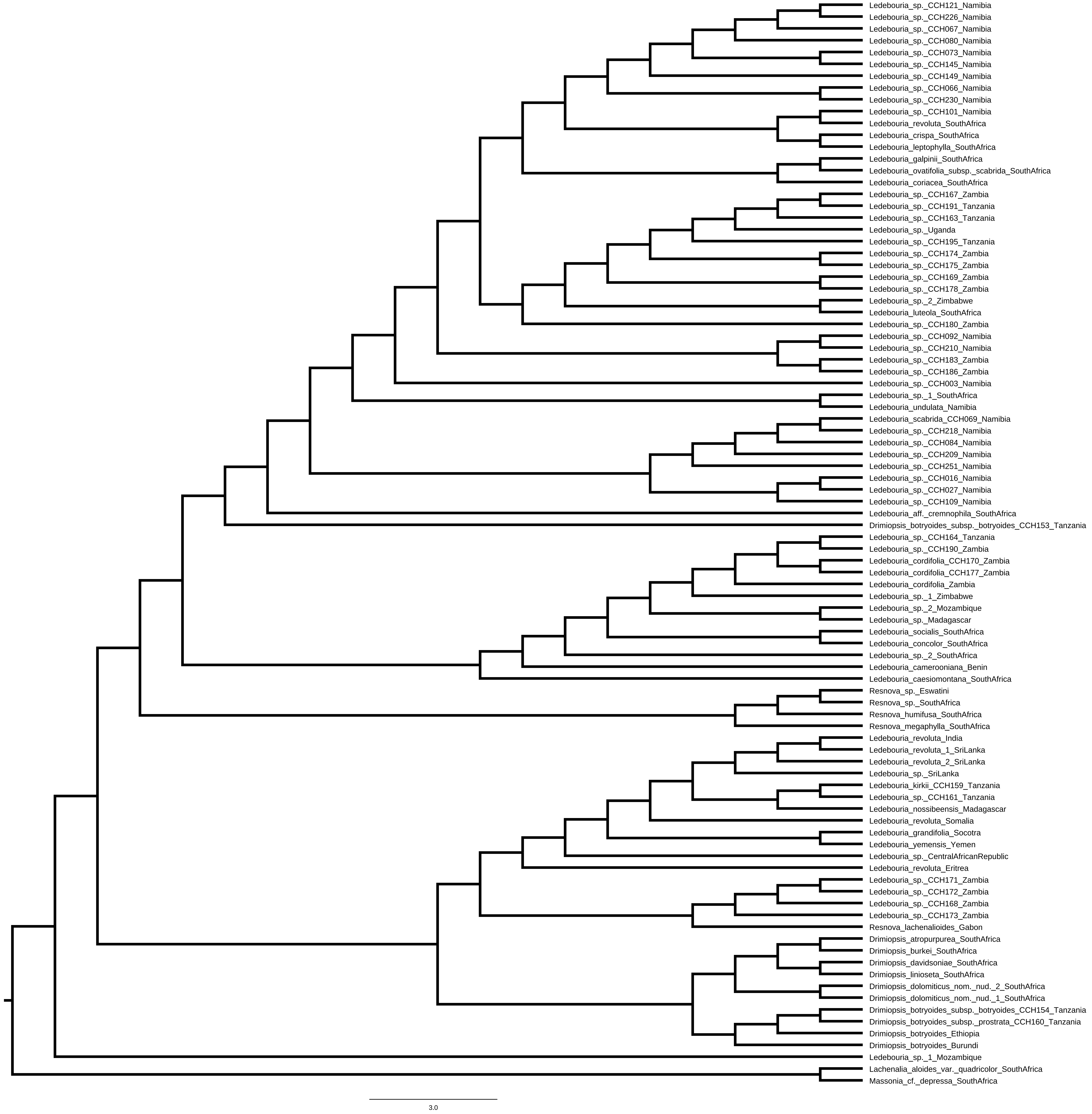

### Fig. S11

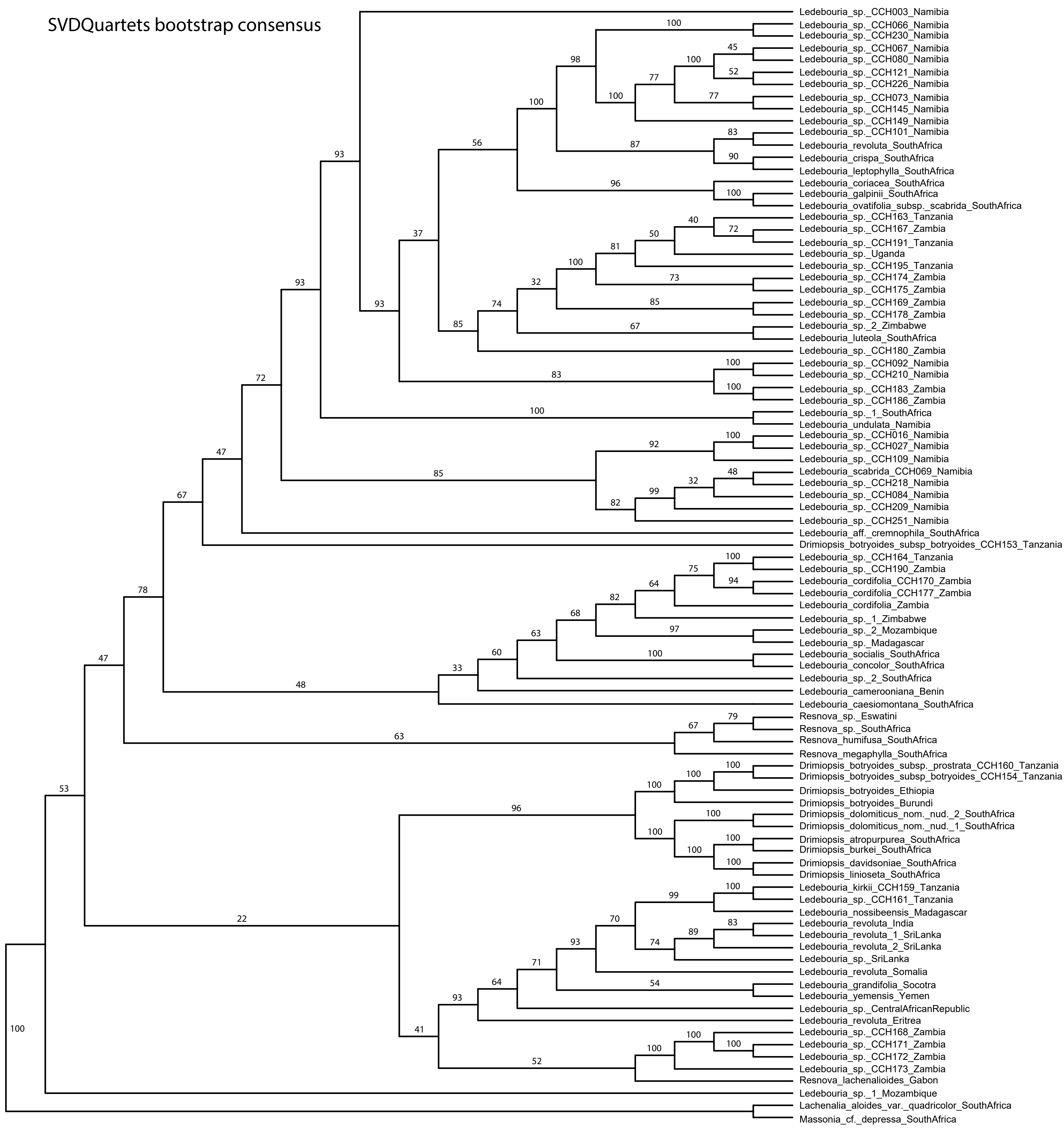

### Fig. S12

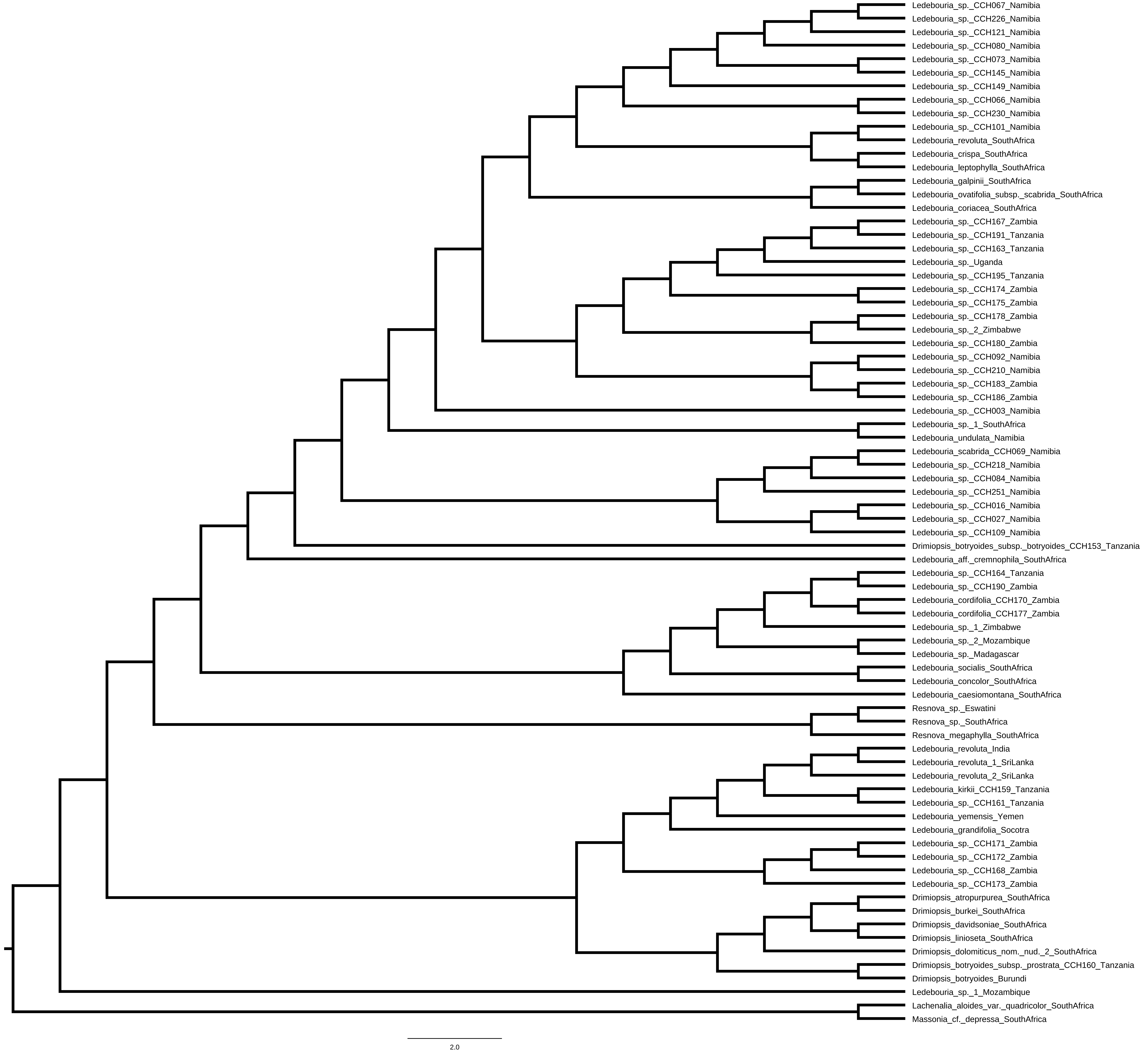

### Fig. S13

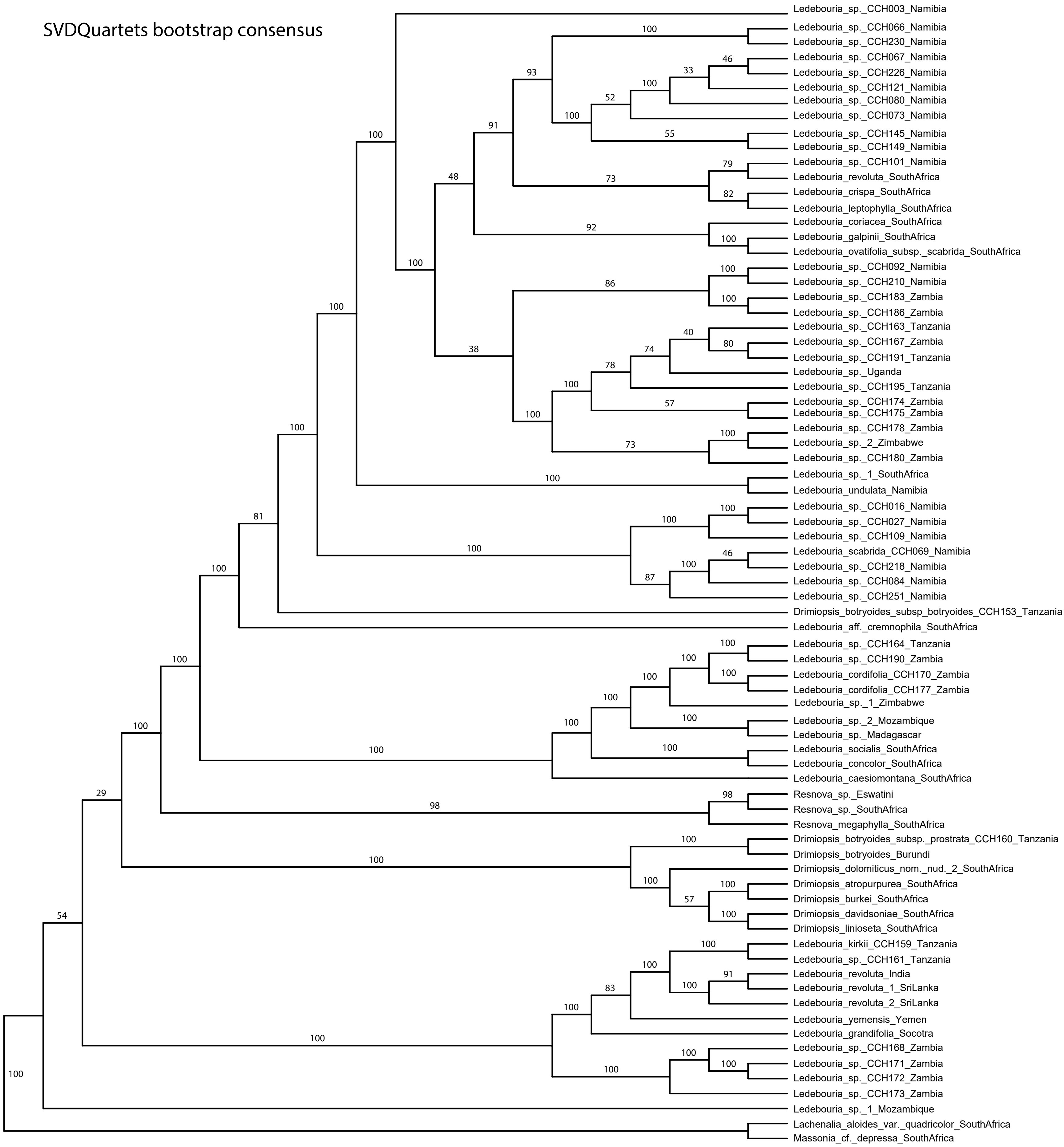

### Fig. S14

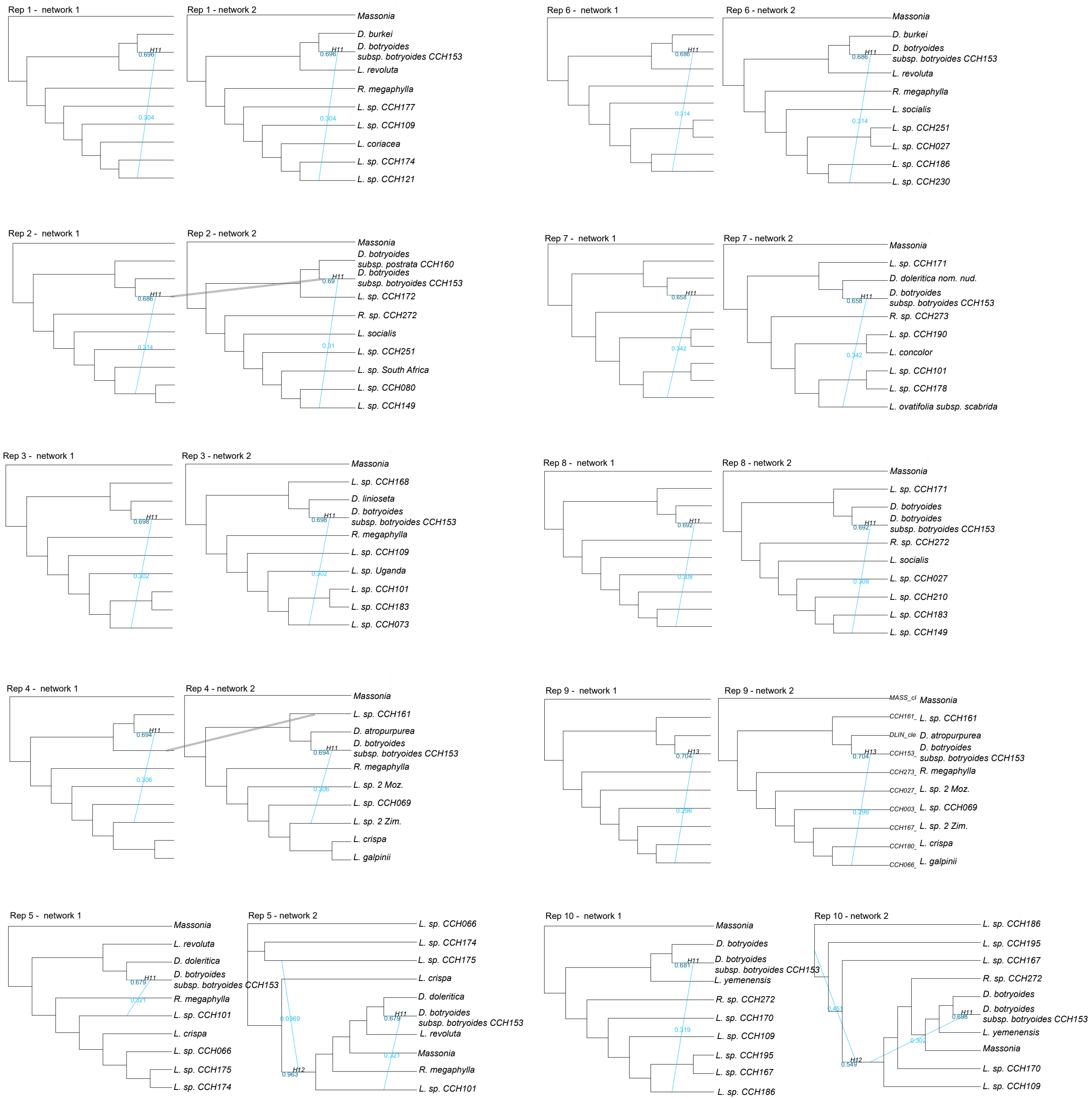

### Fig. S15

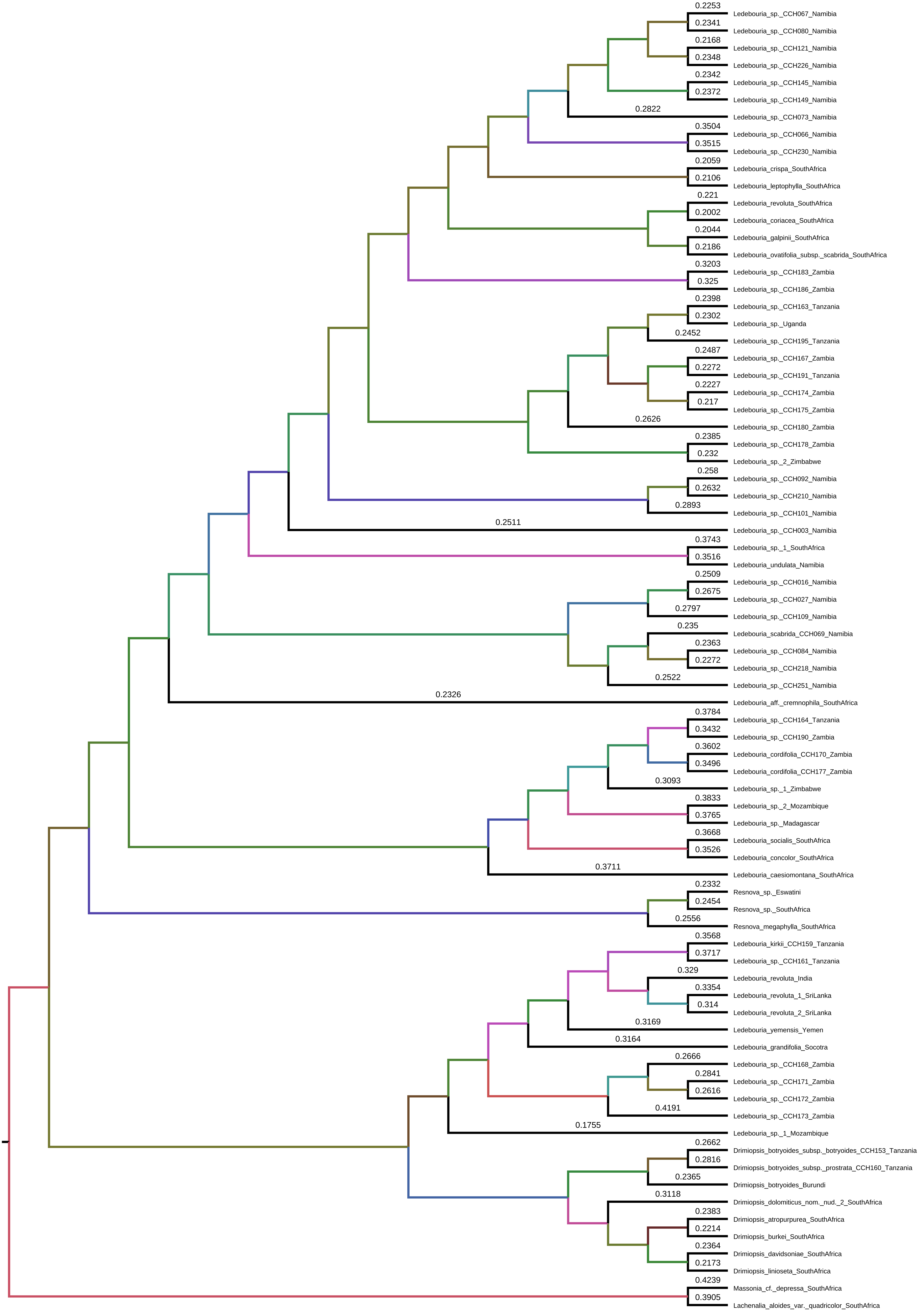

### Fig. S16

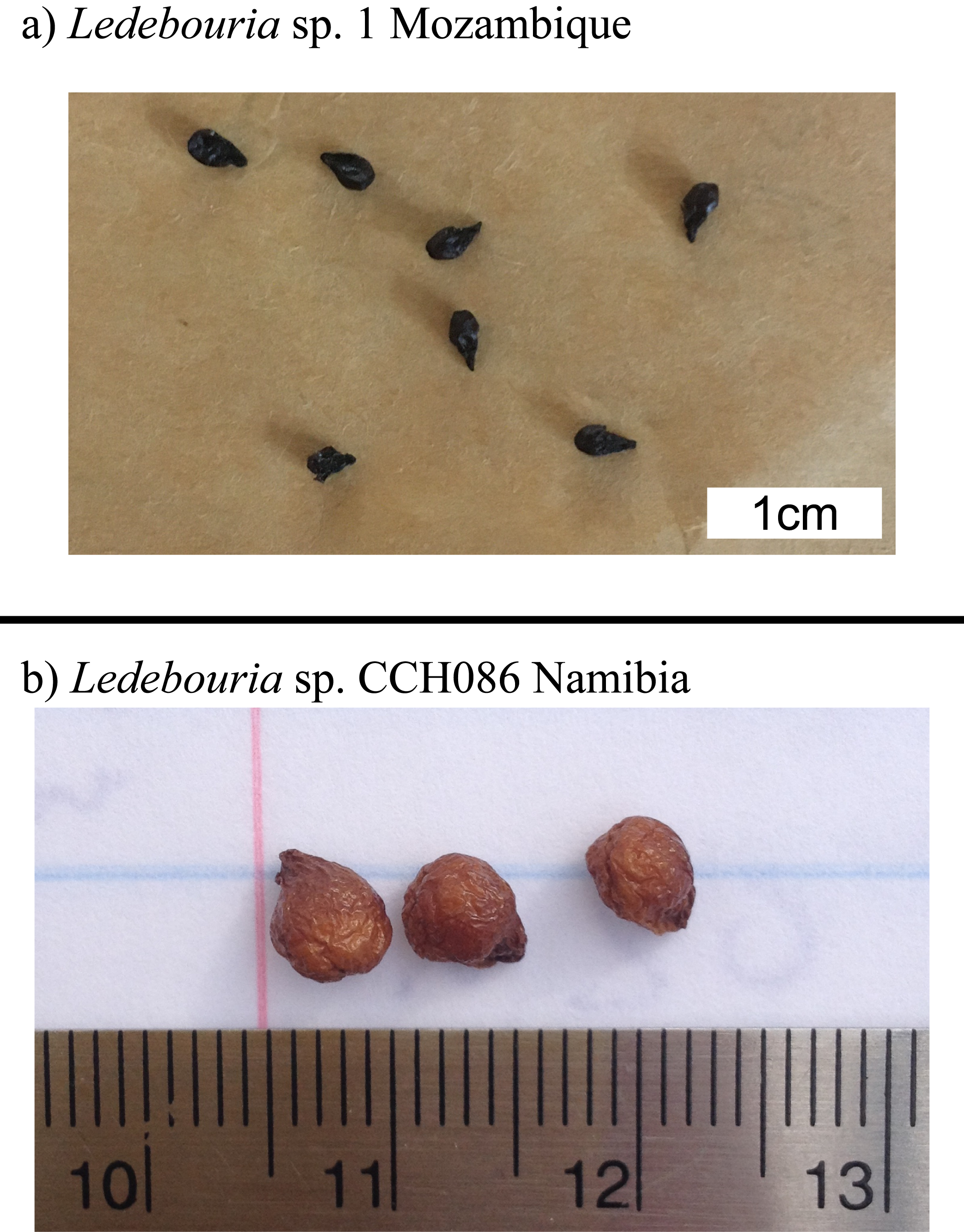
