## Supplementary material for "Peeling Back the Layers: First Phylogenomic Insights into the Ledebouriinae (Scilloideae, Asparagaceae)": Table S1

| genus | specific epithet | collection information | country of origin | material type |
|---|---|---|---|---|
| Drimiopsis | botryoides subsp. botryoides | CCH153 | Tanzania | living |
| Drimiopsis | botryoides subsp. botryoides | CCH154 | Tanzania | living |
| Drimiopsis | botryoides subsp. prostrata | CCH160 | Tanzania | living |
| Drimiopsis | doleritica nom. nud. | CCH241 | South Africa | living |
| Drimiopsis | atropurpurea | CCH278 | South Africa | living |
| Drimiopsis | burkei | CCH238 | South Africa | living |
| Drimiopsis | davidsoniae | CCH270 | South Africa | living |
| Drimiopsis | dolomiticus nom. nud. | CCH241 | South Africa | living |
| Drimiopsis | linioseta | CCH277 | South Africa | living |
| Drimiopsis | botryoides | UPS:BOT:V-640008 | Ethiopia | herbarium |
| Drimiopsis | botryoides | K000771648 | Burundi | herbarium |
| Lachenalia | aloides var. quadricolor | CCH279 | South Africa | living |
| Ledebouria | sp. | CCH003 | Namibia | living |
| Ledebouria | sp. | CCH016 | Namibia | living |
| Ledebouria | sp. | CCH027 | Namibia | living |
| Ledebouria | sp. | CCH066 | Namibia | living |
| Ledebouria | sp. | CCH067 | Namibia | living |
| Ledebouria | scabrida | CCH069 | Namibia | living |
| Ledebouria | sp. | CCH073 | Namibia | living |
| Ledebouria | sp. | CCH080 | Namibia | living |
| Ledebouria | sp. | CCH084 | Namibia | living |
| Ledebouria | sp. | CCH092 | Namibia | living |
| Ledebouria | sp. | CCH101 | Namibia | living |
| Ledebouria | sp. | CCH109 | Namibia | living |
| Ledebouria | sp. | CCH121 | Namibia | living |
| Ledebouria | sp. | CCH145 | Namibia | living |
| Ledebouria | sp. | CCH149 | Namibia | living |
| Ledebouria | kirkii | CCH159 | Tanzania | living |
| Ledebouria | sp. | CCH161 | Tanzania | living |
| Ledebouria | sp. | CCH163 | Tanzania | living |
| Ledebouria | sp. | CCH164 | Tanzania | living |
| Ledebouria | sp. | CCH167 | Zambia | living |
| Ledebouria | sp. | CCH168 | Zambia | living |

| | | | | |
|---|---|---|---|---|
| Ledebouria | sp. | CCH169 | Zambia | living |
| Ledebouria | cordifolia | CCH170 | Zambia | living |
| Ledebouria | sp. | CCH171 | Zambia | living |
| Ledebouria | sp. | CCH172 | Zambia | living |
| Ledebouria | sp. | CCH173 | Zambia | living |
| Ledebouria | sp. | CCH174 | Zambia | living |
| Ledebouria | sp. | CCH175 | Zambia | living |
| Ledebouria | cordifolia | CCH177 | Zambia | living |
| Ledebouria | sp. | CCH178 | Zambia | living |
| Ledebouria | sp. | CCH180 | Zambia | living |
| Ledebouria | sp. | CCH183 | Zambia | living |
| Ledebouria | sp. | CCH186 | Zambia | living |
| Ledebouria | sp. | CCH190 | Zambia | living |
| Ledebouria | sp. | CCH191 | Tanzania | living |
| Ledebouria | sp. | CCH195 | Tanzania | living |
| Ledebouria | sp. | CCH209 | Namibia | living |
| Ledebouria | sp. | CCH210 | Namibia | living |
| Ledebouria | sp. | CCH218 | Namibia | living |
| Ledebouria | sp. | CCH226 | Namibia | living |
| Ledebouria | sp. | CCH230 | Namibia | living |
| Ledebouria | socialis | CCH244 | South Africa | living |
| Ledebouria | concolor | CCH246 | South Africa | living |
| Ledebouria | sp. | CCH251 | Namibia | living |
| Ledebouria | revoluta | CCH262 | South Africa | living |
| Ledebouria | caesiomontana | CCH280 | South Africa | living |
| Ledebouria | coriacea | CCH254/HBG104939 | South Africa | living |
| Ledebouria | aff. cremnophila | CCH281 | South Africa | living |
| Ledebouria | crispa | HBG81213 | South Africa | living |
| Ledebouria | galpinii | HBG93169 | South Africa | living |
| Ledebouria | grandifolia | DH98058 | Socotra | living |
| Ledebouria | sp. 1 | CCH282 | South Africa | living |
| Ledebouria | leptophylla | CCH283 | South Africa | living |
| Ledebouria | sp. 1 | E-3-15 | Mozambique | living |
| Ledebouria | sp. 1 | B-3-15 | Zimbabwe | living |

| | | | | |
|---|---|---|---|---|
| Ledebouria | sp. 2 | C-3-15 | Mozambique | living |
| Ledebouria | sp. 2 | R-3-15 | Zimbabwe | living |
| Ledebouria | yemensis | | Yemen | living |
| Ledebouria | ovatifolia subsp. scabrida | CCH276 | South Africa | living |
| Ledebouria | sp. | | Uganda | living |
| Ledebouria | undulata | CCH284 | Namibia | living |
| Ledebouria | revoluta 1 | S09-27618 | Sri Lanka | herbarium |
| Ledebouria | revoluta | S12-29117 | Somalia | herbarium |
| Ledebouria | revoluta | UPS:BOT:V-057079 | Eritrea | herbarium |
| Ledebouria | camerooniana | MO6564566 | Benin | herbarium |
| Ledebouria | revoluta 2 | S09-27293 | Sri Lanka | herbarium |
| Ledebouria | sp. 2 | MO3814873 | South Africa | herbarium |
| Ledebouria | sp. | MO3288337 | Central Afrian Republic | herbarium |
| Ledebouria | luteola | MO6488867 | South Africa | herbarium |
| Ledebouria | cordifolia | MO6227178 | Zambia | herbarium |
| Ledebouria | nossibeensis | MO6111318 | Madagascar | herbarium |
| Ledebouria | sp. | MO6291239 | Madagascar | herbarium |
| Ledebouria | revoluta | K:TB12038 | India | herbarium |
| Massonia | cf. depressa | CCH285 | South Africa | living |
| Resnova | sp. | CCH272 | Eswatini | living |
| Resnova | sp. | CCH273 | Mkuze, South Africa | living |
| Resnova | megaphylla | CCH286 | South Africa | living |
| Resnova | lachenalioides | K000771641 | Gabon | herbarium |
| Resnova | humifusa | K000771755 | South Africa | herbarium |
